# A chemogenetic light-activatable reactive oxygen species generator for precise cell manipulation

**DOI:** 10.64898/2026.11.12.766124

**Authors:** Hela Benaissa, Stephanie Board, Sabine Tricot, Karim Ounoughi, Cyprien Velmir, Isabelle Aujard, Thomas Le Saux, Ludovic Jullien, Anna Campalans, Arnaud Gautier

**Author notes:** These authors contributed equally (Hela Benaissa, Stephanie Board).

## Abstract

The ability to generate reactive oxygen species (ROS) through illumination of photosensitizers provides a powerful means to manipulate cellular function with high spatial and temporal resolution. Here, we present a chemogenetic light-activatable ROS generator based on the FAST technology. By leveraging the heavy atom effect, we engineered chromophores – called ROSgen – that can both fluoresce and produce ROS when bound to FAST variants and exposed to light. Photosensitization is strictly dependent on the addition of the ROSgen, allowing for on-demand activated and targeted photomanipulation. We demonstrate that this chemogenetic light-activatable ROS generator enables to selectively perturb the function of organelles such as mitochondria, inactivate proteins or induce localized oxidative DNA damage, demonstrating its versatility as molecular tool for manipulating cellular processes with high spatial and temporal control.

## Introduction

Reactive oxygen species (ROS) are naturally produced within cells and are carefully regulated to maintain physiological balance. Excessive intracellular ROS can, however, cause oxidative damage and perturb cells^1^. The ability to induce ROS production precisely, both in space and time, has become invaluable in biological research. By employing specially designed photosensitizers that generate ROS only when exposed to specific wavelength of light, researchers can control or perturb cells with remarkable accuracy, targeting specific proteins for inactivation, ablating single cells, or perturbing specific signaling pathways^2–8^.

Photosensitizers operate through intricate photochemical mechanisms. Upon light absorption, they are promoted into an excited singlet state that may return to the ground state via fluorescence or undergo intersystem crossing to form an excited triplet state. Although this transition is formally spin-forbidden, many photosensitizers are engineered or naturally predisposed to facilitate this transition efficiently. Once in the triplet excited state, the photosensitizer can interact with molecular oxygen – which naturally exists in a triplet ground state – to produce singlet oxygen or other ROS.

Genetically-encoded photosensitizers, such as KillerRed^5^ and miniSOG^9^, enables to achieve precise localized ROS generation in living cells. Genetic targeting allows the selective ablation of specific cells in a population or the selective inactivation of a proximal target. The emergence of chemogenetic photosensitizers, which combines exogenous chromophores with protein scaffolds, has further enabled the creation of systems that can be activated on-demand for ROS production^2,4,10^. For instance, fluorogen-activating proteins (FAP) were engineered into targeted and activated photosensitizers (TAP) by incorporating iodine into malachite green dye, thus promoting ROS production when exposed to far-red light^4^. The incorporation of heavy atoms as iodine enhances spin-orbit coupling and promotes efficient intersystem-crossing, making these chromophores efficient light-activated ROS sources when bound to FAP. Similarly, rhodamine-based HaloTag ligands were chemically modified with heavy atoms to efficiently generate ROS upon green light illumination^2^.

Fluorescent genetically targeted photosensitizers have also been shown to be powerful probes for correlative light and electron microscopy. By generating highly localized ROS, they can precisely convert 3,3’-diaminobenzidine into a highly localized polymeric precipitate, which can be stained with osmium tetroxide for visualization by electron microscopy^2,9^. Furthermore, the unique properties of photosensitizers have inspired innovative light-controlled approaches for proximity labeling, enabling the mapping of subcellular transcriptome^11^, the probing of protein-protein interactions^12^ or the profiling of subcellular proteome^13^ with unprecedented spatial resolution.

In this study, we leverage the modularity of the FAST (Fluorescence-Activating and absorption-Shifting Tag) technology to develop a targetable and activatable chemogenetic photosensitizer capable of generating ROS on demand upon blue light illumination. FAST and its variants are small 14 kDa protein tags binding selectively and non-covalently fluorogenic dyes, producing bright fluorescent assemblies^14–20^. By incorporating heavy atoms such as bromine and iodine, we took advantage of the heavy atom effect to increase ROS production^21^. Like their non-halogenated counterparts, these modified chromophores dissipate light energy non-radiatively in solution and in cells, remaining non-fluorescent and inert. Only upon binding to FAST, do they become fluorescent and produce ROS. We call these activatable chromophores “ROSgens” or “ROSgenic chromophores” to highlight their unique capability to light-induce ROS exclusively upon FAST binding. This system enables precise, on-demand control of photoinduced reactions by simply adding ROSgenic chromophores. Here we show that FAST-based chemogenetic ROS generator enables to perturb cell function with light in a controlled manner, opening new ways for the precise control of biological processes.

## Results & Discussion

### Design and characterization of ROSgens

To create FAST-based ROS generators, we leveraged the molecular tunability of 4-hydroxybenzylidene rhodanine (HBR) derivatives. These push-pull chromophores – composed of an electron-donating phenol conjugated to an electron-withdrawing rhodanine – exhibit high fluorescence efficiency when they adopt a planar conformation in the cavity of prototypical FAST but are otherwise non-fluorescent in solution^14,16,20,22^. The introduction of heavy atoms – such as bromine or iodine – to chromophores is known to significantly increase their ability to generate ROS by enhancing the rate of intersystem crossing^4,10^. We thus synthesized a set of HBR analogs, modifying the ortho positions of the phenol with bromine and iodine, to explore and enhance their photosensitizing properties (**Figure 1a,b**).

**Figure 1.**
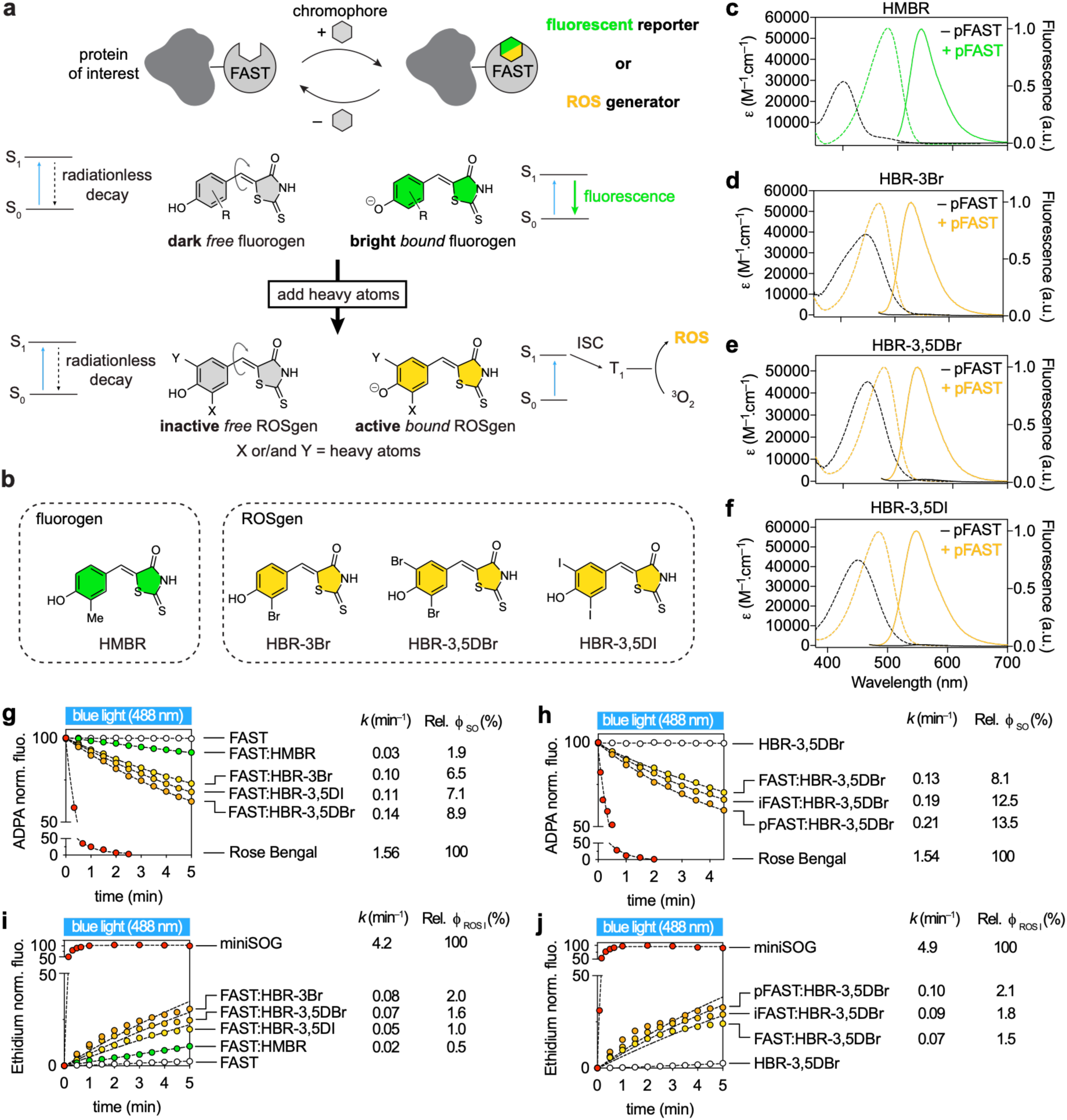
Light-activatable chemogenetic reactive oxygen species generator. **a** pFAST can form ROS generator upon binding to heavy-atom-substituted HBR analogs. **b** Structures of the chromophores used in this study. **c-f** Absorption (dashed lines) and fluorescence (solid lines) spectra of (**c**) HMBR, (**d**) HBR-3Br, (**e**) HBR-3,5DBr and (**f**) HBR-3,5DI when free in solution (dark lines) and bound to pFAST (colored lines). Spectra were recorded using chromophore concentrations at 3 μM and pFAST at 40 μM in pH 7.4 phosphate buffer saline at 25°C. **g,h** Solutions of ADPA and various photosensitizers were irradiated with 488 nm light (90 mW.cm^−2^). ADPA fluorescence was measured after different exposure times (Ex 374 nm, Em 384 – 550 nm) (see also **Supplementary Figures 2 and 3**). Rose Bengal (RB) was used as a standard and control of singlet oxygen generation. Data were fitted using one phase exponential decay equation. The absorption at 488 nm was identical in all samples (0.055 ± 10%), enabling direct comparison of the kinetics and estimation of the relative singlet oxygen quantum yields (ϕ_SO_). **i,j** Solutions of dihydroethidium (HE) and various photosensitizers were irradiated with 488 nm light (90 mW.cm^−2^). Ethidium fluorescence was measured after different exposure times (Ex 525 nm, Em 600 – 750 nm) (see also **Supplementary Figures 5 and 6**). Values were normalized by the maximal fluorescent intensities obtained with miniSOG and data were fitted using one phase exponential association equation. The absorption at 488 nm was identical in all samples (0.055 ± 10%), enabling direct comparison of the kinetics and estimation of the relative Type I ROS quantum yields (ϕ_ROS I_).

We observed that, as regular HBR derivatives, these heavy-atom-substituted HBR derivatives are non-fluorescent in solution, but form tight fluorescent assemblies with the variants FAST^14^, iFAST^22^ and pFAST^20^ (**Figure 1c-f** and **Table 1**). The wavelengths of absorption and emission, as well as the molar absorptivity of these bimolecular assemblies, were similar to those observed with the standard fluorogen HMBR (**Figure 1c-f** and **Table 1**). However, we observed a notable decrease in fluorescence quantum yield: about 40% lower for HBR-3,5DI and nearly 70% for both HBR-3,5DBr and HBR-3Br (**Table 1**). This reduction implies that a part of the absorbed light energy is dissipated through other deexcitation pathways, in agreement with enhanced intersystem crossing.

**Table 1.** Properties of FAST variants with heavy-atom-substituted HBR analogs.

| **Mutant** | **Fluorogen** | **λ<sub>abs</sub> (nm)** | **Δλ<sub>abs</sub> (nm)** | **λ<sub>em</sub> (nm)** | **ε (mM<sup>-1</sup>cm<sup>-1</sup>)** | **ϕ<sub>F</sub>** | **Molecular brightness** | ***K*<sub>D</sub> (µM)** |
|---|---|---|---|---|---|---|---|---|
| FAST | HMBR (ref<sup>16</sup>) | 481 | 79 | 540 | 44 | 0.23 | 10,000 | 0.13 |
|  | HBR-3,5DI | 484 | 33 | 547 | 51 | 0.15 | 7,600 | 0.11 |
|  | HBR-3,5DBr | 474 | 20 | 536 | 45 | 0.07 | 3,100 | 0.04 |
|  | HBR-3Br | 469 | 30 | 529 | 51 | 0.07 | 3,500 | 0.07 |
| iFAST | HMBR (ref<sup>22</sup>) | 480 | 78 | 541 | 41 | 0.22 | 9,000 | 0.07 |
|  | HBR-3,5DI | 481 | 30 | 544 | 55 | 0.13 | 7,200 | 0.07 |
|  | HBR-3,5DBr | 473 | 20 | 536 | 54 | 0.06 | 3,200 | 0.02 |
|  | HBR-3Br | 469 | 29 | 530 | 48 | 0.06 | 3,000 | 0.02 |
| pFAST | HMBR (ref<sup>20</sup>) | 481 | 79 | 542 | 54 | 0.23 | 13,000 | 0.01 |
|  | HBR-3,5DI | 486 | 34 | 548 | 56 | 0.14 | 7,900 | 0.01 |
|  | HBR-3,5DBr | 475 | 30 | 536 | 50 | 0.06 | 3,000 | 0.02 |
|  | HBR-3Br | 470 | 23 | 530 | 54 | 0.07 | 3,800 | 0.005 |

Abbreviations are as follows: λ<sub>abs</sub> wavelength of maximal absorption; Δλ<sub>abs</sub> = λ<sub>abs,bound</sub> - λ<sub>abs,unbound</sub> absorption red-shift upon chromophore binding; λ<sub>em</sub> wavelength of maximal emission; ε, molar absorptivity at λ<sub>abs</sub> (standard error is typically 10%); ϕ<sub>F</sub> fluorescence quantum yield; molecular brightness = ϕ × ε ; *K*<sub>D</sub> thermodynamic dissociation constant.

To evaluate the light-induced ROS generation capabilities of heavy-atom-substituted HBR analogs bound to FAST variants, we conducted experiments in vitro using anthracene-9,10-dipropionic acid (ADPA), a widely used singlet oxygen probe^23^. Singlet oxygen reacts with anthracene derivatives to form oxidized products^24^, thus monitoring ADPA photooxidation provides an indirect measure of singlet oxygen production^4,9^. It is however important to note that this photobleaching reaction is not entirely specific to singlet oxygen, as anthracene can also be oxidized via electron transfer process, which may lead to an overestimation of the quantum yield of singlet oxygen generation^25–27^. Bearing this caveat in mind, we used ADPA photobleaching to characterize various assemblies formed between FAST variants and heavy-atom-substituted HBR analogs. During these assays, samples were continuously illuminated with 488 nm light, and the fluorescence intensity of ADPA was measured over time to estimate the efficiency of singlet oxygen generation (**Figure 1g,h, Supplementary Figures 1 and 2**). As a standard, Rose Bengal (RB), a well-characterized chemical photosensitizer^28^, was included for comparison. To ensure that ROS generation did not result in substantial destruction of the photosensitizer itself, we also monitored the fluorescence of the assemblies under identical illumination conditions, confirming minimal photobleaching throughout the experiment (**Supplementary Figure 3**). Comparative analysis revealed that FAST assemblies with heavy-atom substituted HBR analogs displayed significantly enhanced singlet oxygen generation relative to the standard FAST:HMBR assembly: we observed increases of 240% for FAST:HBR-3,5DI, 270% for FAST:HBR-3Br and 360% for FAST:HBR-3,5DBr. Moreover, the assemblies iFAST:HBR-3,5DBr and pFAST:HBR-3,5DBr were 50% and 60% more efficient, respectively, than FAST:HBR-3,5DBr, suggesting that the protein environment can further modulate the efficiency of singlet oxygen production for a given chromophore. The top-performing combination, pFAST:HBR-3,5DBr, achieved 13.5% of the singlet oxygen generation efficiency of Rose Bengal (**Figure 1g,h**).

To further assess the generation of other ROS, we employed the dihydroethidium (HE) probe which is selectively oxidized by species such as superoxide and hydrogen peroxide^29^. In its reduced form, HE emits blue fluorescence, but upon oxidation, it is converted to ethidium that fluoresces strongly at 610 – 630 nm. Since singlet oxygen does not oxidize HE^26,30^, this probe serves as a useful indicator of type I ROS production. During our experiment, photosensitizer solutions containing HE were illuminated with 488 nm light and the formation of ethidium was tracked over time, allowing us to estimate the efficiency of type I ROS generation (**Figure 1i,j, Supplementary Figures 4 and 5**). As a benchmark, we used miniSOG, reported to generate efficiently superoxide and hydrogen peroxide^26,27^. We observed that none of the ROSgenic complexes matched the efficiency of miniSOG in oxidizing HE (**Figure 1i,j**): after 5 minutes of irradiation, with pFAST:HBR-3,5DBr and FAST:HBR-3Br, only about 30% of HE was oxidized, a level achieved within just 5 s with miniSOG. However, although our chemogenetic photosensitizers generate type I ROS with modest efficiency, we observed that incorporating heavy atoms did enhance their ability to produce type I ROS, with pFAST being a slightly superior scaffold compared to FAST and iFAST (**Figure 1i,j**).

### Selective imaging in live cells

Next, we tested the use of heavy-atom substituted HBR analogs for the labeling of pFAST in live cells. Preliminary tests revealed that treating mammalian cells with ROSgenic dyes at concentrations used for labeling (5 μM) for up to 24 h caused minimal effects on cell viability (**Supplementary Figure 6**). Although less bright than with regular HBR derivatives, pFAST assemblies with heavy-atom substituted HBR derivatives were sufficiently fluorescent to visualize targeted proteins in live cells using confocal microscopy. By applying low excitation power and brief exposures, we successfully imaged HeLa cells expressing pFAST in the cytoplasm, mitochondria and at the inner plasma membrane, without inducing any observable morphology change (**Supplementary Figure 7**).

### Targeted light-induced perturbation of mitochondria

Directing ROS generation to specific subcellular compartments, such as mitochondria, cell membranes, or nuclei, can cause substantial changes. ROS are known to cause DNA damage, oxidize membrane lipids, and trigger various cell death pathways, especially within mitochondria^31^. To assess our system’s capacity for localized ROS production and cellular perturbation, we expressed pFAST in mitochondria (using a mito-pFAST fusion) in HeLa cells. Cells were treated with HBR-3,5DBr, and illuminated with low-intensity 488 nm light for 10 min. This illumination was intentionally kept gentle to avoid non-specific phototoxicity. Mitochondrial morphology was tracked using co-expressed mito-mCherry and 560 nm light excitation. In cells treated with HBR-3,5DBr, 488 nm illumination specifically induced the formation of globular and donut-shaped (toroidal) mitochondria (**Figure 2a**). Such morphology changes – known to be linked to mitochondria dysfunction – are thought to arise from osmotic stress and oxidative damage, and have previously been associated with loss of mitochondrial membrane potential^32,33^. In contrast, control cells treated with HMBR showed no such change (**Figure 2b**), suggesting that the observed mitochondrial alterations were due to light-triggered generation of ROS by pFAST:HBR-3,5DBr. Additional controls using mito-miniSOG under the same illumination conditions led to more pronounced signs of cell damage, including mitochondria fragmentation and cell shrinkage (**Figure 2c**). Note that miniSOG fluorescence rapidly photobleached during illumination, suggesting self-inactivation, whereas pFAST:HBR-3,5DBr maintained its fluorescence throughout the experiment. The gentler ROS generation ability of pFAST:HBR-3,5DBr enabled here to finely tune photooxidation, allowing us to visualize subtle changes in mitochondria morphology, changes inaccessible with a stronger ROS generator like MiniSOG. This result suggests that such gentler ROS generator can provide advantages for nuanced targeted modulation of cellular processes and fine tuning of cell physiology without causing irreversible damage.

**Figure 2.**
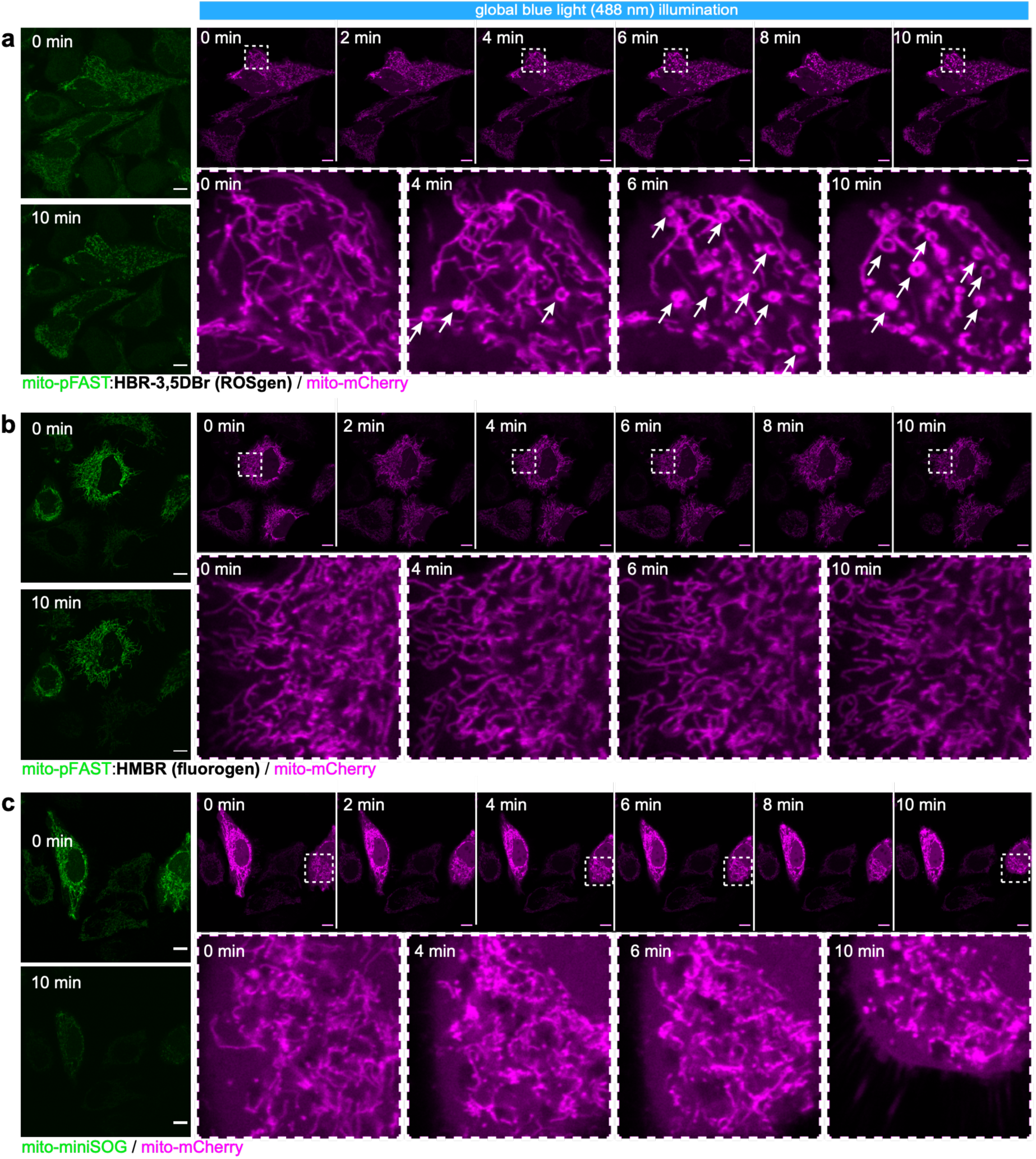
Illumination of mitochondria-localized pFAST:ROSgen can induce selective mitochondria morphology change. Confocal micrographs of live HeLa cells co-expressing **(a,b)** mito-pFAST or **(c)** mito-miniSOG and **(a,b,c)** mito-mCherry (mito sequence = mitochondrial targeting sequence from the subunit VIII of human cytochrome C oxidase). HeLa cells were illuminated with 488 nm light (light intensity 13.7 kW/cm^2^, pixel dwell time 4.9 μs) for 10 minutes (1 scan / 5 sec). The fluorescence of (**a**,**b**) mito-pFAST in presence of (**a**) 10 μM HBR-3,5DBr and (**b**) 10 μM HMBR, and of (**c**) mito-miniSOG were recorded before and after irradiation. Mitochondria morphology was monitored with the fluorescence of co-expressed mito-mCherry. Experiments were repeated three times with similar results. Scale bars, 10 μm.

Next, we tested whether light-induced ROS generation causes mitochondrial membrane depolarization. We pre-labeled mitochondria with tetramethylrhodamine ethyl ester (TMRE), a dye that accumulates in polarized mitochondria proportionally to their membrane potential. Upon membrane depolarization, TMRE is released out of mitochondria, resulting in a loss of fluorescence. We treated cells expressing mito-pFAST with HBR-3,5DBr and 10 nM TMRE, then illuminated them with 488 nm light for 2 min while imaging TMRE using 561 nm excitation. After 2 min of light exposure, we observed an average loss of 75 % in TMRE fluorescence in pFAST-expressing cells (**Figure 3a**), while non-transfected cells retained their fluorescence, indicating that membrane depolarization was specifically induced by the FAST system. In control experiments using HMBR instead of HBR-3,5DBr, only a 20 % decrease of fluorescence was observed under identical conditions (**Figure 3b**), showing the superior efficiency of pFAST:HBR-3,5DBr complex in generating ROS and inducing phototriggered depolarization of mitochondrial membrane.

**Figure 3.**
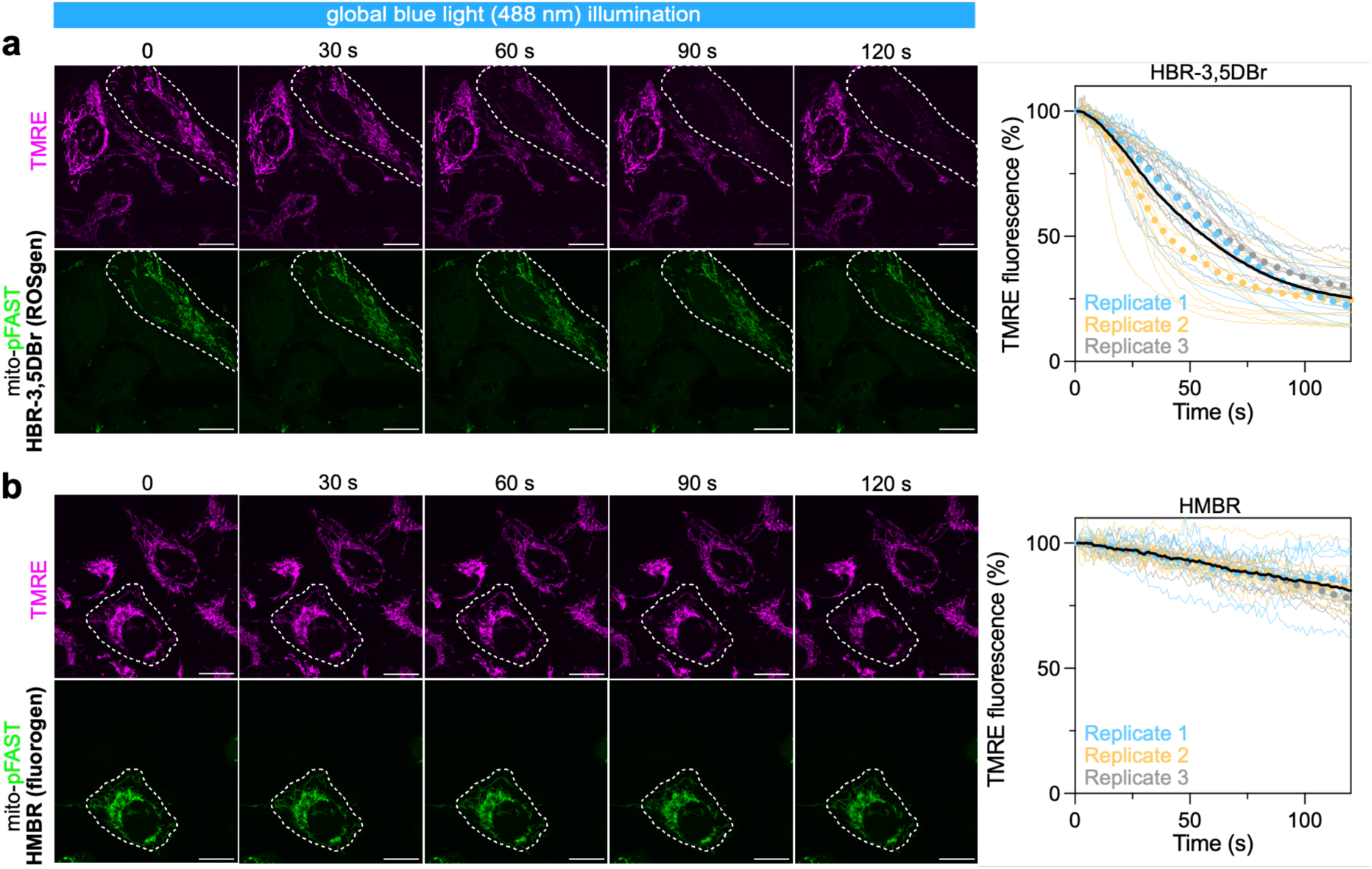
Illumination of mitochondria-localized pFAST:ROSgen can induce selective mitochondrial membrane depolarization. **a,b** Representative two-color time-lapse confocal imaging of HeLa cells expressing mito-pFAST (green), stained with TMRE (magenta) to monitor mitochondrial membrane potential, and treated with (**a**) HBR-3,5DBr or (**b**) HMBR. Cells were illuminated across the full field with 488 nm light for 2 min (1 scan/s). Dotted outlines mark pFAST-positive cells. The graphs show the evolution of TMRE fluorescence from 30 pFAST-positive cells across three independent experiments. Each cell is color-coded according to the biological replicate it came from. The dotted lines represent the mean value of each biological replicate, while the black line represents the mean of the biological replicates. Scale bars, 20 μm.

By applying localized irradiation to a subregion of the mitochondria network in cells expressing mito-pFAST treated with HBR-3,5DBr, we were able to selectively trigger membrane depolarization in individual cells within seconds (**Figure 4a,b**). We observed that although damages are induced only locally, membrane depolarization occurs on the entire mitochondrial network in agreement with the electric connectivity of mitochondrial network as previously reported^34,35^. Control experiments replacing HBR-3,5DBr by HMBR or omitting pFAST did not show such global membrane depolarization (**Figure 4c-f**), in agreement with selective ROS generation upon illumination of pFAST:HBR-3,5DBr. This set of experiments demonstrates the high spatial and temporal resolution of the approach.

**Figure 4.**
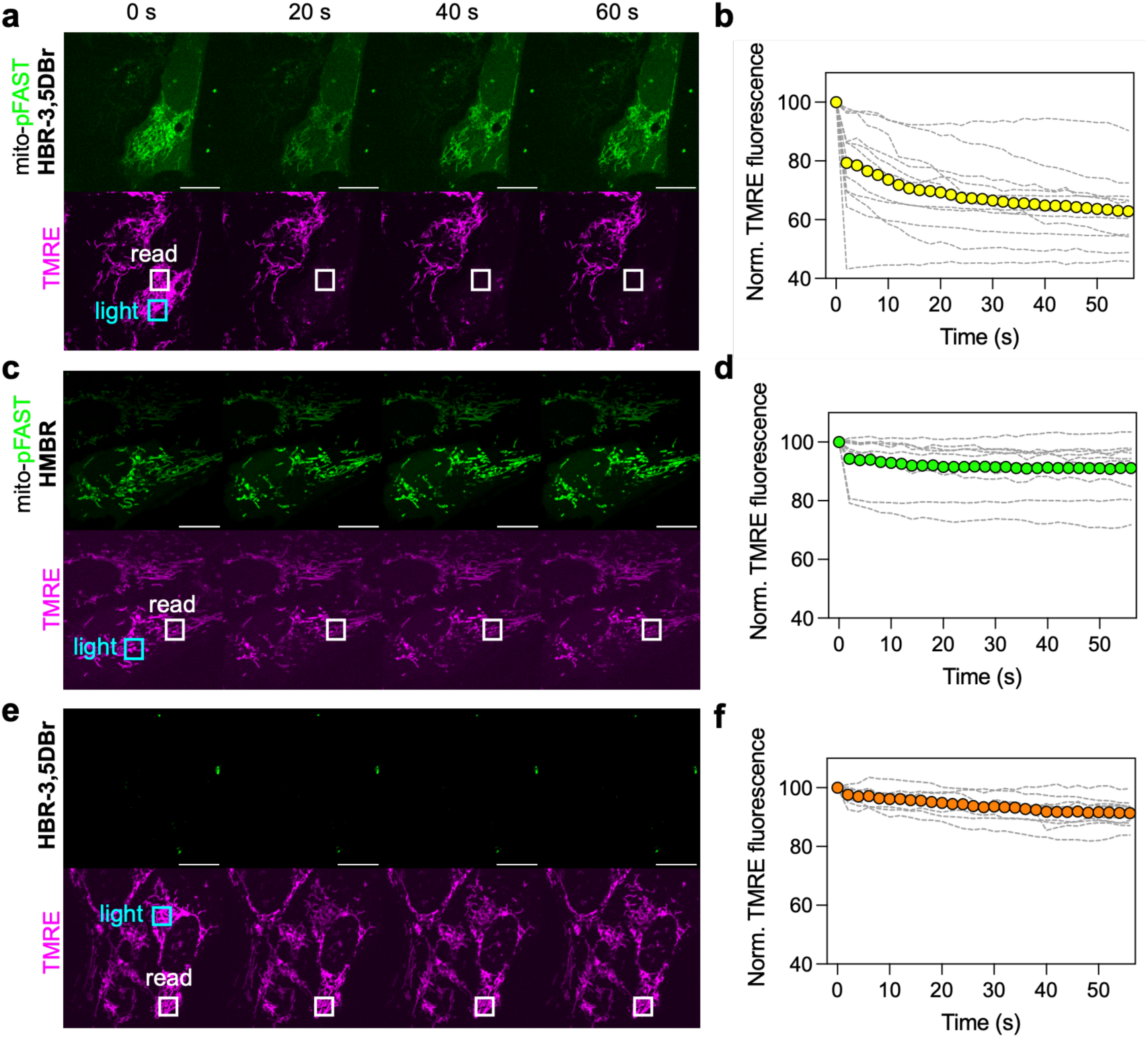
Local illumination of mitochondria-localized pFAST:ROSgen can induce selective mitochondrial membrane depolarization. **a,c,e** Representative two-color time-lapse confocal imaging of U2OS cells expressing mito-pFAST (green) (**a,c**) or not (**e**), stained with TMRE (magenta) to monitor mitochondrial membrane potential, and treated with (**a,e**) HBR-3,5DBr or (**c**) HMBR. Cells were illuminated locally with 488 nm light (blue square) for 10 sec (1 scan/s). TMRE fluorescence was measured over time in the white square (read). **b,d,f** The graphs show the evolution of TMRE fluorescence from n = 12 (**b**), 11 (**d**), 11(**f**) individually irradiated cells. The solid circles represent the mean value of the replicates. Scale bars, 20 μm.

### Chromophore-assisted light inactivation

We next tested the use of FAST:HBR-3,5DBr for chromophore-assisted light inactivation (CALI) of proteins in live cells. CALI leverages targeted ROS generation to selectively inactivate proteins fused to photosensitizers by local photooxidation, offering precise spatial and temporal control over protein function^5^. As a proof-of-concept, we chose as a model protein the δ1 pleckstrin (PLC) homology (PH) domain. This protein localizes at the inner leaflet of the plasma membrane interacting with membrane phospholipids. Prior studies have demonstrated that ROS-mediated CALI can disrupt this interaction, causing the protein to translocate from the membrane to the cytosol^5^. We constructed a triple fusion protein mCherry-pFAST-(PH)_2_, combining the fluorescent protein mCherry for imaging, pFAST for light-induced ROS production, and two repeats of the PH domain for efficient membrane localization. After expressing this construct in HEK293T cells and incubation with HBR-3,5DBr, we illuminated cells with 488 nm light while monitoring mCherry localization at 561 nm. Upon illumination, we observed a pronounced increase in cytosolic mCherry fluorescence (**Figure 5a**), indicating successful CALI-induced release of the protein from the membrane to the cytosol. Control experiments with HMBR displayed minimal relocation (**Figure 5b**), consistent with the reduced ROS-generating capacity of pFAST:HMBR. Similarly, cells lacking either the chromophore or pFAST showed negligible changes (**Figure 5c,d**), confirming the specificity and efficacy of pFAST:HBR-3,5DBr for CALI-mediated protein inactivation in living cells.

**Figure 5.**
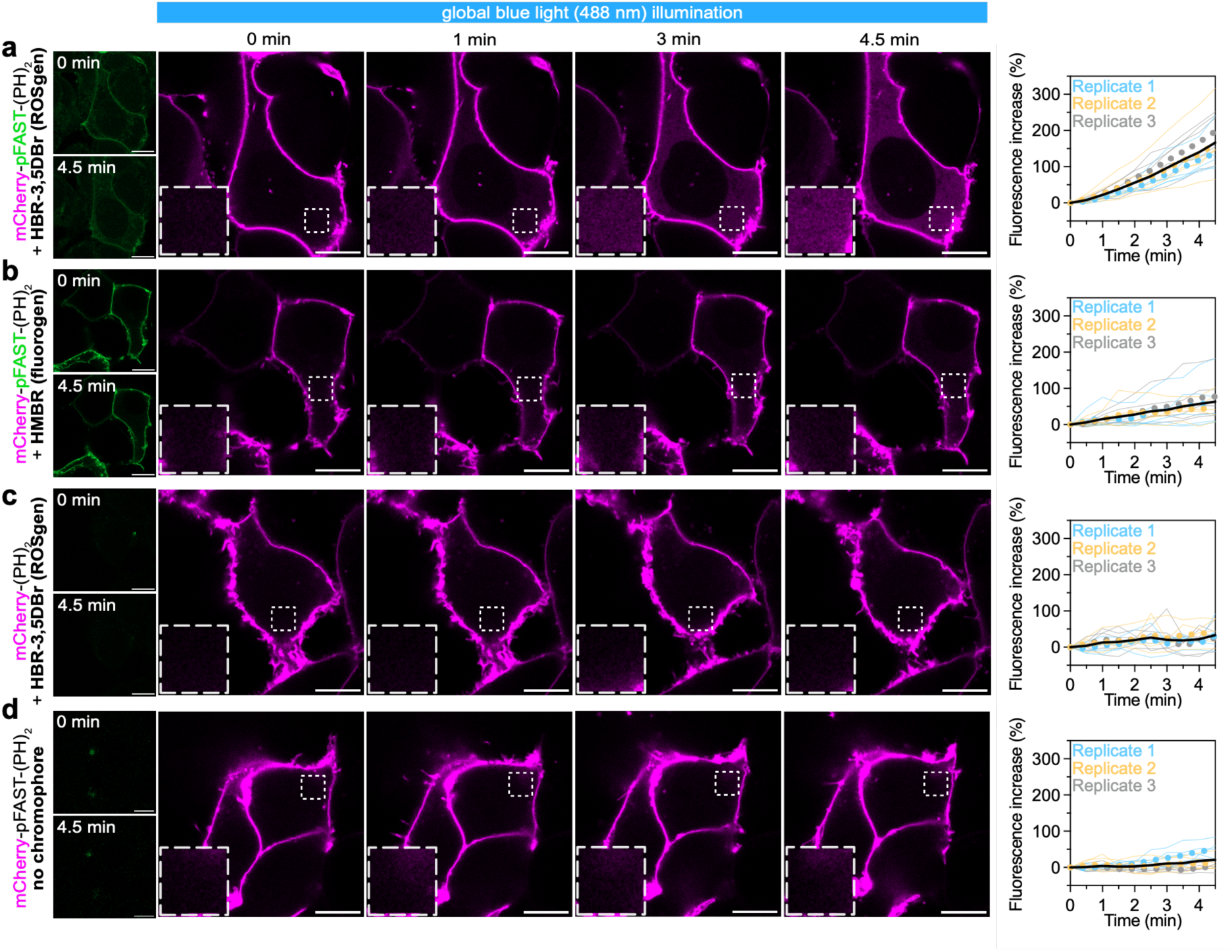
pFAST:ROSgen enables chromophore-assisted light inactivation (CALI). Representative time-lapse confocal imaging of HEK cells expressing either mCherry-pFAST-(PH)_2_ (**a**,**b**,**d**) or mCherry-(PH)_2_ alone (**c**) during 488 nm illumination in the presence of HBR-3,5DBr (**a**,**c**), HMBR (**b**), or no chromophore (**d**). Images show redistribution of mCherry fluorescence over 4.5 min. Boxed regions indicate areas of interest and are shown magnified. Scale bars, 10 μm. The graphs show quantification of cytoplasmic mCherry fluorescence intensity over time of 15 cells from three independent experiments. Each cell is color-coded according to the biological replicate it came from. The dotted lines represent the mean value of each biological replicate, while the black line represents the mean of the biological replicates.

### Targeted light-induced oxidative DNA damage

We then explore the potential of FAST-based ROS generator to induce oxidative DNA damage in the nucleus of living cells upon light activation. ROS readily target DNA, with guanine being particularly vulnerable due to its low redox potential^36^. Oxidation of guanine results in the formation of 8-oxo-7,8-dihydroguanine (8-oxoG), a mutagenic lesion linked to cancer and age-related disorders, as it can cause G:C to T:A transversions^37^. The accumulation of mutations is limited in cells thanks to the repair of the lesion by the base excision repair (BER) pathway, initiated by the enzyme OGG1 that specifically recognizes and excise the 8-oxoG^38^. To investigate light-induced oxidative DNA damage using pFAST-based ROS generator, we used H2B-pFAST fusion to specifically localize pFAST to the nuclear DNA. Upon illumination with 488 nm light, cells expressing H2B-pFAST showed a marked and selective increase in 8-oxoG lesions (**Figure 6a,b**), detected by immunostaining with a specific anti-8-oxoG antibody. Microirradiation within the nucleus of H2B-pFAST-expressing cells led to localized 8-oxoG lesions (**Figure 6c,d**), demonstrating that local illumination of pFAST-based ROS-generator can induce localized DNA photooxidation. Overall, this set of experiment shows that pFAST-based ROS-generator can enable to oxidize DNA with very high spatial and temporal resolution, opening new perspective for the study of oxidative DNA damages.

**Figure 6.**
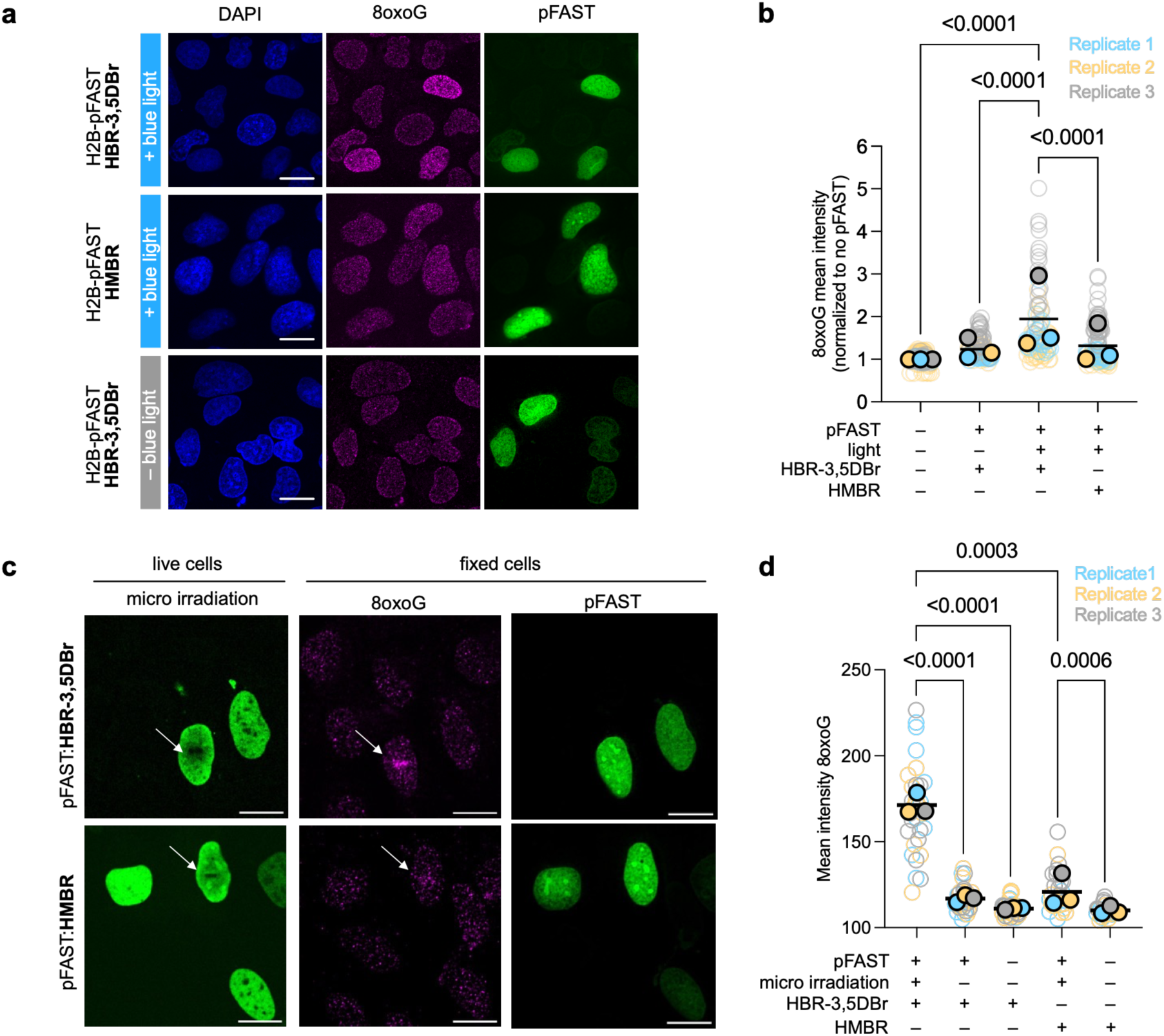
pFAST:ROSgen enables DNA photooxidation. **a,b** U2OS cells expressing H2B-pFAST were treated with HBR-3,5DBr or HMBR and illuminated with blue light. Cells were then fixed, immunostained with an antibody against 8-oxoG and labeled with HMBR to visualize pFAST and DAPI to stain DNA. Control experiment with no light illumination was also performed. Representative micrographs are shown. Scale bars, 20 μm. The graph shows the quantification of 8-oxoG from three independent experiments. Each cell is color-coded according to the biological replicate it came from. The solid circles represent the mean value of each biological replicate, while the black line represents the mean of the biological replicates. Statistical analysis was performed using a Kruskal–Wallis test and a Dunn’s test. **c,d** U2OS cells expressing H2B-pFAST were treated with HBR-3,5DBr or HMBR and illuminated locally with blue light (white arrow indicates the area of illumination). Cells were then fixed, immunostained with an antibody against 8-oxoG and labeled with HMBR to visualize pFAST. Control experiment with no light illumination was also performed. Representative micrographs are shown. Scale bars, 20 μm. The graph shows the quantification of 8-oxoG from three independent experiments. Each cell is color-coded according to the biological replicate it came from. The solid circles represent the mean value of each biological replicate, while the black line represents the mean of the biological replicates. Statistical analysis was performed using a Kruskal–Wallis test and a Dunn’s test.

### Conclusion

The ability to generate ROS with light opens a multitude of possibilities for manipulating and studying biological systems, including targeted organelle disruption, selective protein inactivation, cellular ablation, correlative light and electronic microscopy or proximity labeling. In this work, we show that introducing heavy atoms in HBR analogs allows the generation of pFAST-based chemogenetic photosensitizer with enhanced ROS-generating capabilities.

Photosensitization is initiated by the strict addition of the ROSgenic chromophores, allowing for on-demand activated and targeted photomanipulation. Our results show that this pFAST-based ROS generator can enable to selectively perturb the function of organelles such as mitochondria, inactivate proteins through CALI or induce localized DNA damage, demonstrating its versatility as molecular tool for manipulating cellular processes with spatial and temporal precision. Compared to other classical photosensitizers, such as miniSOG, our pFAST-based ROS generator enables more gentle ROS production, ideal for nuanced, targeted modulation of cellular processes – offering a powerful means for finely tuning cell physiology without causing irreversible damage. Interestingly, the dual capacity of pFAST – to bind either a fluorogen or a ROSgen – offers remarkable versatility as it is possible to switch between imaging and photomanipulation by simply exchanging the chromophore, greatly broadening the experimental potential of this single molecular platform.

## DATA AVAILABILITY

The data supporting the findings of this study are available within the article and supplementary information, and are available from the corresponding authors upon reasonable request. Source data are provided with this paper.

## ACKNOWLEDGMENTS

We thank the microscopy facility of the Institut de Biologie Paris Seine of Sorbonne University, and more particularly France Lam and Chloé Chaumeton for their assistance. This work has been supported by the European Research Council (ERC-2016-CoG-724705 FLUOSWITCH), the Agence Nationale de la Recherche (ANR-21-CE11-0010-02 FluNanoTrack to AG and ANR-23-CE12-0025 OXIREPTRA to AC) and the Institut Universitaire de France.

## AUTHOR CONTRIBUTIONS

H.B., S.B. and A.G. designed the overall project and wrote the paper with the help of the other authors. H.B., S.B., L.J., A.C. and A.G. designed the experiments. H.B., S.B., S.T., K.O., C.V., I.A., A.C. performed the experiments. H.B., S.B., S.T., K.O., C.V., I.A, L.J., A.C. and A.G. analyzed the experiments.

## COMPETING INTERESTS

The authors declare the following competing financial interest: A.G. and L.J. are co-founders and hold equity in Twinkle Bioscience/The Twinkle Factory, a company commercializing the FAST technologies. The other authors declare no competing interests.

## MATERIALS & METHODS

### Organic synthesis

Commercially available reagents were used as obtained. ^1^H and ^13^C NMR spectra were recorded at 300 K on a Bruker AM 300 spectrometer; chemical shifts are reported in ppm with protonated solvent as internal reference ^1^H, CHD_2_SOCD_3_ in CD_3_SOCD_3_ 2.49 ppm; coupling constants J are given in Hz. Mass spectra high resolution were performed by the Service de Spectrométrie de Masse de l’Institut de Chimie Organique et Analytique (Orléans). The preparation of HMBR (4-hydroxy-3-methylbenzylidene rhodanine) was previously described^14^.

### HBR-3Br (4-hydroxy-3-bromobenzylidene)-3-phenyl-2-thioxothiazolidin-4-one)

Solid 4-hydroxy-3-bromobenzaldehyde (241 mg, 1.2 mmol) and rhodanine (100 mg, 0.75 mmol) were mixed and the mixture was brought to 170°C. It first fused and then solidified. After cooling to 50-60°C, ethanol was added and the mixture was heated at reflux until complete dissolution. Then an excess of water was added and a precipitate appeared. The precipitate was filtered and dried over P_2_O_5_. HBR-3Br was obtained as a yellow powder (60 %, 142 mg). ^1^H NMR (300 MHz, DMSO-*d6*, δ): 13.73 (s, 1H), 11.26 (s, 1H), 7.79 (s, 1H), 7.54 (s, 1H), 7.43 (d, *J* = 8 Hz, 1H), 7.10 (d, *J* = 8 Hz, 1H); ^13^C NMR (75 MHz, DMSO-*d6*, δ): 195.3, 169.4, 156.6, 136.0, 130.9, 130.7, 125.6, 122.7, 112.1, 110.3; HRMS (ESI): m/z 315.9096 (calcd mass for C_10_H_7_BrNO_2_S_2_ [M+H]^−^: 315.9096) (in accordance with previously described analysis^39,40^).

### HBR-3,5DBr (4-hydroxy-3,5-dibromobenzylidene)-3-phenyl-2-thioxothiazolidin-4-one)

Solid 3,5-dibromo-4-hydroxybenzaldehyde (252 mg, 0.9 mmol), rhodanine (100 mg, 0.75 mmol) and piperidine (22 μL, 0.2 mmol) were mixed in ethanol (3 mL) and stirred at reflux temperature for 24h. After cooling, the precipitate formed during the reaction was collected by filtration, washed with cold ethanol and dried under reduced pressure. HBR-3,5DBr was obtained as an orange powder (78%, 232 mg). ^1^H NMR (300 MHz, DMSO-d6, δ): 7.77 (s, 2H), 7.55 (s, 1H); HRMS (ESI): m/z 393.8202 (calcd mass for C_10_H_6_Br_2_NO_2_S_2_ [M+H]^−^: 393.8201) (in accordance with previously described analysis^41^).

### HBR-3,5DI

(4-hydroxy-3,5-diiodobenzylidene)-3-phenyl-2-thioxothiazolidin-4-one). Solid 3,5-diiodo-4-hydroxybenzaldehyde (342 mg, 0.9 mmol), rhodanine (100mg, 0.75 mmol), and piperidine (22 μL, 0.2 mmol) were mixed in ethanol (3 mL) and stirred at reflux temperature for 24h. After cooling, the precipitate formed during the reaction was collected by filtration, washed with cold ethanol and dried under reduced pressure. HBR-3,5DI was obtained as a red powder (57%, 209 mg). ^1^H NMR (300 MHz, DMSO-*d6*, δ): 7.95 (s, 2H), 7.49 (s, 1H); HRMS (ESI): m/z 489.7917 (calcd mass for C_10_H_6_I_2_NO_2_S_2_ [M+H]^-^: 489.7923) (in accordance with previously described analysis^41^).

### Molecular cloning

Synthetic oligonucleotides used for cloning were purchased from Integrated DNA Technology. PCR reactions were performed with Q5 polymerase (New England Biolabs) following manufacturer guidelines. PCR products and restriction digest fragments were loaded to 1-2% agarose gel for 45 minutes at 100 volts in TAE buffer and purified using QIAquick Gel Extraction Kit PCR purification kit (Qiagen). Cloning was performed using restriction digest and Gibson assembly methods. Restriction enzymes were purchased by NEB or Thermofisher scientific and isothermal assembly (Gibson assembly) were performed using mix prepared according to previously described protocols^1^. Hybridization of short oligos were performed using a ramp down method in a thermocycler starting at 95°C reducing 1°C per cycle for 70 cycles. Constructs were ligated using T4 ligase following manufacturer guidelines (NEB). All products were transformed in DH10 beta (NEB) and DB3.1 (Thermofisher) E. coli strains using either electroporation or chemically competent methods. Small-scale isolation of plasmid DNA was done using QIAprep miniprep kit (Qiagen) from 3 mL of overnight culture supplemented with appropriate antibiotics. Large-scale isolation of plasmid DNA was done using the QIAprep maxiprep kit (Qiagen) from 150 mL of overnight culture supplemented with appropriate antibiotics. All constructs were cloned into a common vector with a pBR322 origin of replication site and a CMV promoter sequence. All plasmid sequences were confirmed by Sanger sequencing with appropriate sequencing primers or whole plasmid sequencing (Eurofins genomics). The plasmids allowing the mammalian expression of pFAST, or pFAST fused to the mitochondrial targeting sequence from the subunit VIII of human cytochrome C oxidase (Mito-pFAST), to the inner plasma membrane targeting sequence lyn11 (Lyn11-pFAST) and to the histone H2B (H2B-pFAST) under the control of a CMV promoter were previously described^42^.

The plasmid pAG675 for bacterial expression of miniSOG with an N-Terminal His-tag under the control of a T7 promoter (pAG675) was constructed by replacing the sequence coding for FAST in the plasmid pAG87^14^ using isothermal Gibson assembly. The plasmid pAG935, pAG1633 and pAG1634 for the mammalian expression of mito-MiniSOG, mCherry-pFAST-(PH)_2_ and mCherry-(PH)_2_ were constructed by isothermal Gibson assembly. See **Supplementary Table 2** for ORF sequences.

### Characterization of the chemogenetic photosensitizer complexes

#### Chemicals

Anthracene-9,10-dipropionic acid disodium salt (ADPA) was obtained from Coger, dihydroethidium (HE) (≥ 95%) and Rose Bengal (RB) (95%) were obtained from Sigma-Aldrich and used as received.

#### Protein expression

pAG675 and the plasmids driving *E. coli* expression of FAST^14^, iFAST^22^, pFAST^42^ with an N-terminal His-tag were transformed in Rosetta (DE3) pLys *E. coli* (New England Biolabs). Cells were grown at 37°C in Lysogen Broth (LB) medium supplemented with 50 μg/mL kanamycin and 34 μg/mL chloramphenicol to OD_600nm_ 0.6. Expression was induced overnight at 16°C by adding isopropyl-β-D-1-thiogalactopyranoside (IPTG) to a final concentration of 1 mM. Cells were harvested by centrifugation (4,300 × g for 20 min at 4°C) and frozen.

#### Protein purification

The cell pellet was resuspended in lysis buffer (PBS supplemented with 2.5 mM MgCl_2_, 1mM of protease inhibitor PhenylMethaneSulfonyl Fluoride PMSF, 0.025 mg/mL of DNAse, pH 7.4) and sonicated (5 min, 20 % of amplitude) on ice. The lysate was incubated for 2-3 hours on ice to allow DNA digestion by DNAse. Cellular fragments were removed by centrifugation (9,000 × g for 1 h at 4°C). The supernatant was incubated overnight at 4°C by gentle agitation with pre-washed Ni-NTA agarose beads in PBS buffer complemented with 20 mM of imidazole. Beads were washed with 10 volumes of PBS complemented with 20 mM of imidazole and with 5 volumes of PBS complemented with 40 mM of imidazole. His-tagged proteins were eluted with 5 volumes of PBS complemented with 0.5 M of imidazole. The buffer was exchanged to PBS (0.05 M phosphate buffer, 0.150 M NaCl) using PD-10 or PD-MidiTrap G-25 desalting columns (GE Healthcare). Purity of the proteins were evaluated using SDS-PAGE electrophoresis stained with Coomassie blue.

#### Physico-chemical measurements

Steady state UV-Vis absorption spectra were recorded using a Cary 300 UV-Vis spectrometer (Agilent Technologies), equipped with a Versa20 Peltier-based temperature-controlled cuvette chamber (Quantum Northwest) and fluorescence spectra were recorded using a LPS 220 spectrofluorometer (PTI, Monmouth Junction, NJ), equipped with a TLC50TM Legacy/PTI Peltier-based temperature-controlled cuvette chamber (Quantum Northwest). Fluorescence quantum yield measurements were determined as previously described using FAST:HMBR as a reference^14,43^. Reciprocal dilution with protein solution was used so as to keep the protein concentration constant at 40 μM while diluting the fluorogen solution. Absorption coefficients were determined by forward titration of fluorogens into a 40 μM protein solution. Thermodynamic dissociation constants were determined as previously described^14,43^ using a Spark 10M plate reader (Tecan) and fitting data in Prism 6 to a one-site specific binding model.

#### Reactive Oxygen species quantification

Steady state UV-Vis absorption spectra were recorded using a Cary 300 UV-Vis spectrometer (Agilent Technologies), equipped with a Versa20 Peltier-based temperature-controlled cuvette chamber (Quantum Northwest) and fluorescence spectra were recorded using a LPS 220 spectrofluorometer (PTI, Monmouth Junction, NJ), equipped with a TLC50TM Legacy/PTI Peltier-based temperature-controlled cuvette chamber (Quantum Northwest). For ADPA photobleaching, HE photooxidation and photobleaching experiments, the samples were irradiated with a current-controlled blue color Light Emitting Diodes LED (M470L4, Thorlabs, NJ) (filtered at λ = 488 ± 20 nm) aligned in a 90° degree angle with respect to the photomultiplier detector of the spectrofluorometer. We relied on the photoswitching kinetics of a photochemically well-characterized RSFP (Dronpa2 at pH 7.4) for measuring light intensities of the LED light (as previously described^44,45^). All experiments were performed in 3 mm^2^ ultra-micro cells quartz cuvettes to avoid any effect from diffusion. For ADPA photobleaching measurements, all samples were incubated with 50 μM of ADPA and fluorescence was monitored at 374 nm excitation and 384 – 550 nm emission as a function of the duration of 488 nm light exposure (90 mW.cm^−2^). For HE oxidation measurements, all samples were incubated with 50 μM of HE and increase of fluorescent products was monitored at 525 nm excitation and 600 – 750 nm emission as a function of the duration of 488 nm light (90 mW.cm^−2^).

### Mammalian cell culture

All reagents were purchased from Thermofisher Scientific unless stated otherwise. HEK 293T cells (ATCC CRL-3216) were cultured in Dulbecco’s modified Eagle’s medium (DMEM) supplemented with high Glucose, L-glutamine, Phenol Red, Sodium Pyruvate and 10% (vol/vol) fetal calf serum, at 37 °C in a 5% CO_2_ atmosphere. HeLa cells (ATCC CCL-2) were cultured in modified Eagle’s medium (MEM) supplemented with phenol red, 1 × non-essential amino acids, 1 × sodium pyruvate, and 10% (vol/vol) fetal calf serum at 37 °C in a 5% CO_2_ atmosphere. U2OS (HTB-96) cells were cultured in McCoy media supplemented with 10% (vol/vol) fetal calf serum at 37 °C in a 5% CO_2_ atmosphere. For imaging, cells were seeded in μDish 35 mm glass bottom dish IBIDI (Clinisciences) coated with poly-L-lysine. Cells were seeded 18 to 24 h prior to transfection at a concentration of 120,000 HEK 293T cells or 60,000 HeLa cells, or 100,000 U2OS cells per ibidi dish. Cells were transiently transfected with 1 μg total plasmid DNA allowing for mammalian expression under the control of a CMV promoter. Transfection was achieved using genejuice (Merck) and serum free media OptiMEM according to the manufacturer’s protocol 24 h prior to microscopy imaging.

### Cell viability assay

HeLa cells were treated with MEM media containing the appropriate ROSgen at the indicated concentrations for the indicated times. The cell viability was evaluated by fluorescence microscopy using the LIVE/DEAD® viability/cytotoxicity assay kit (Molecular Probes, Life Technologies) following the manufacturer’s protocol.

### Immunofluorescence

For immunofluorescence experiments, cells were washed with DMEM and immediately fixed for 5 min at room temperature with cold MeOH/acetone (vol 1:1), air dried and rinsed with PBS. To denaturate DNA, cells were treated with 2N HCl for 45 min at room temperature then washed twice with PBS, neutralized with 50 mM Tris pH 8.8 for 5min at room temperature, washed with PBS and permeabilised at room temperature in PBS 0.1% Triton (×100) for 5 min. Cells were incubated in blocking solution (PBS containing 0.1% Triton 100×, 3% BSA and 1% normal goat serum) at 37°C for 1 h. Cells were incubated with primary antibodies against 8-oxoG (abcam ab48508) diluted at 1/2000 in blocking solution for 1 h at 37°C, washed three times for 5 min in PBS containing 0.1% Triton and further incubated with secondary antibodies Alexa Fluor 594 goat anti Mouse (Fisher, A11005) diluted 1/1000 in blocking solution for 45 min at 37°C. Cells were washed three times for 5 min in PBS containing 0.1% Triton. Nuclear DNA was counterstained with 1 µg/ml 4′,6′-diamidino-2-phenylindole (DAPI).

### Fluorescence microscopy

The confocal micrographs of mammalian cells were acquired on a Leica TCS SP5 confocal laser scanning microscope equipped with a 63× / 1.4 NA oil immersion objective, a Zeiss LSM 980 Laser Scanning Microscope equipped with a plan apochromat 63× / 1.4 NA oil immersion objective and a Nikon Yokogawa CSU-W1 spinning disk microscope equipped with a Plan-Apochromat 60× / 1.42 NA oil immersion objective. Live cells were washed twice with DMEM media (without serum and phenol red) immediately prior to imaging. To the same DMEM media, the fluorogens/ROSgen were diluted at the indicated concentration and added to the cells. Cells were imaged directly without washing. Leica LAS AF software, ZEN software or Matamorph were used to collect the data. The images were analysed with Fiji (Image J) and data processed using GraphPad Prism 7.

## References

1. Schieber, M. & Chandel, N. S. ROS Function in Redox Signaling and Oxidative Stress. Current Biology 24, R453–R462 (2014).

2. Binns, T. C., Ayala, A. X., Grimm, J. B., Tkachuk, A. N., Castillon, G. A., Phan, S., Zhang, L., Brown, T. A., Liu, Z., Adams, S. R., Ellisman, M. H., Koyama, M. & Lavis, L. D. Rational design of bioavailable photosensitizers for manipulation and imaging of biological systems. Cell Chem. Biol. 27, 1063–1072.e7 (2020).

3. Liao, J. C., Roider, J. & Jay, D. G. Chromophore-assisted laser inactivation of proteins is mediated by the photogeneration of free radicals. PNAS 91, 2659–2663 (1994).

4. He, J., Wang, Y., Missinato, M. A., Onuoha, E., Perkins, L. A., Watkins, S. C., St. Croix, C. M., Tsang, M. & Bruchez, M. P. A near-infrared genetically targetable and activatable photosensitizer. Nat Methods 13, 263–268 (2016).

5. Bulina, M. E., Chudakov, D. M., Britanova, O. V., Yanushevich, Y. G., Staroverov, D. B., Chepurnykh, T. V., Merzlyak, E. M., Shkrob, M. A., Lukyanov, S. & Lukyanov, K. A. A genetically encoded photosensitizer. Nat. Biotechnol. 24, 95–99 (2006).

6. Makhijani, K., To, T.-L., Ruiz-González, R., Lafaye, C., Royant, A. & Shu, X. Precision Optogenetic Tool for Selective Single- and Multiple-Cell Ablation in a Live Animal Model System. Cell Chem Biol 24, 110–119 (2017).

7. Bulina, M. E., Lukyanov, K. A., Britanova, O. V., Onichtchouk, D., Lukyanov, S. & Chudakov, D. M. Chromophore-assisted light inactivation (CALI) using the phototoxic fluorescent protein KillerRed. Nature Protocols 1, 947–953 (2006).

8. Riani, Y. D., Matsuda, T., Takemoto, K. & Nagai, T. Green monomeric photosensitizing fluorescent protein for photo-inducible protein inactivation and cell ablation. BMC Biol. 16, 50 (2018).

9. Shu, X., Lev-Ram, V., Deerinck, T. J., Qi, Y., Ramko, E. B., Davidson, M. W., Jin, Y., Ellisman, M. H. & Tsien, R. Y. A genetically encoded tag for correlated light and electron microscopy of intact cells, tissues, and organisms. PLoS Biol. 9, e1001041 (2011).

10. Ren, K., Keshri, P., Wu, R., Sun, Z., Yu, Q., Tian, Q., Zhao, B., Bagheri, Y., Xie, Y. & You, M. A Genetically Encoded RNA Photosensitizer for Targeted Cell Regulation. Angewandte Chemie 132, 22170–22174 (2020).

11. Wang, P., Tang, W., Li, Z., Zou, Z., Zhou, Y., Li, R., Xiong, T., Wang, J. & Zou, P. Mapping spatial transcriptome with light-activated proximity-dependent RNA labeling. Nat. Chem. Biol. 15, 1110–1119 (2019).

12. To, T.-L., Medzihradszky, K. F., Burlingame, A. L., DeGrado, W. F., Jo, H. & Shu, X. Photoactivatable protein labeling by singlet oxygen mediated reactions. Bioorg. Med. Chem. Lett. 26, 3359–3363 (2016).

13. Zheng, F., Yu, C., Zhou, X. & Zou, P. Genetically encoded photocatalytic protein labeling enables spatially-resolved profiling of intracellular proteome. Nat. Commun. 14, 2978 (2023).

14. Plamont, M.-A., Billon-Denis, E., Maurin, S., Gauron, C., Pimenta, F. M., Specht, C. G., Shi, J., Quérard, J., Pan, B., Rossignol, J., Moncoq, K., Morellet, N., Volovitch, M., Lescop, E., Chen, Y., Triller, A., Vriz, S., Le Saux, T., Jullien, L. & Gautier, A. Small fluorescence-activating and absorption-shifting tag for tunable protein imaging in vivo. Proc. Natl. Acad. Sci. U. S. A. 113, 497–502 (2016).

15. Li, C., Mourton, A., Plamont, M. A., Rodrigues, V., Aujard, I., Volovitch, M., Le Saux, T., Perez, F., Vriz, S., Jullien, L., Joliot, A. & Gautier, A. Fluorogenic Probing of Membrane Protein Trafficking. Bioconjug. Chem. 29, 1823–1828 (2018).

16. Li, C., Plamont, M.-A., Sladitschek, H. L., Rodrigues, V., Aujard, I., Neveu, P., Le Saux, T., Jullien, L. & Gautier, A. Dynamic multicolor protein labeling in living cells. Chemical Science 8, 5598–5605 (2017).

17. Li, C., Tebo, A. G., Thauvin, M., Plamont, M.-A., Volovitch, M., Morin, X., Vriz, S. & Gautier, A. A far-red emitting fluorescent chemogenetic reporter for in vivo molecular imaging. Angew. Chem. Int. Ed Engl. 59, 17917–17923 (2020).

18. El Hajji, L., Bunel, B., Joliot, O., Li, C., Tebo, A. G., Rampon, C., Volovitch, M., Fischer, E., Pietrancosta, N., Perez, F., Morin, X., Vriz, S. & Gautier, A. A tunable and versatile chemogenetic near-infrared fluorescent reporter. Nat. Commun. 16, 2594 (2025).

19. Gautier, A. Fluorescence-activating and absorption-shifting tags for advanced imaging and biosensing. Acc. Chem. Res. 55, 3125–3135 (2022).

20. Benaissa, H., Ounoughi, K., Aujard, I., Fischer, E., Goïame, R., Nguyen, J., Tebo, A. G., Li, C., Le Saux, T., Bertolin, G., Tramier, M., Danglot, L., Pietrancosta, N., Morin, X., Jullien, L. & Gautier, A. Engineering of a fluorescent chemogenetic reporter with tunable color for advanced live-cell imaging. Nat. Commun. 12, 6989 (2021).

21. Koziar, J. C. & Cowan, D. O. Photochemical heavy-atom effects. Acc. Chem. Res. 11, 334–341 (1978).

22. Tebo, A. G., Pimenta, F. M., Zhang, Y. & Gautier, A. Improved chemical-genetic fluorescent markers for live cell microscopy. Biochemistry 57, 5648–5653 (2018).

23. Hoebeke, M. & Damoiseau, X. Determination of the singlet oxygen quantum yield of bacteriochlorin a: a comparative study in phosphate buffer and aqueous dispersion of dimiristoyl-L-alpha-phosphatidylcholine liposomes. Photochem Photobiol Sci 1, 283–287 (2002).

24. Lindig, B. A., Rodgers, M. A. J. & Schaap, A. P. Determination of the lifetime of singlet oxygen in water-d2 using 9,10-anthracenedipropionic acid, a water-soluble probe. J. Am. Chem. Soc. 102, 5590–5593 (1980).

25. Kotani, H., Ohkubo, K. & Fukuzumi, S. Photocatalytic Oxygenation of Anthracenes and Olefins with Dioxygen via Selective Radical Coupling Using 9-Mesityl-10-methylacridinium Ion as an Effective Electron-Transfer Photocatalyst. J. Am. Chem. Soc. 126, 15999–16006 (2004).

26. Pimenta, F. M., Jensen, R. L., Breitenbach, T., Etzerodt, M. & Ogilby, P. R. Oxygen-Dependent Photochemistry and Photophysics of “MiniSOG,” a Protein-Encased Flavin. Photochemistry and Photobiology 89, 1116–1126 (2013).

27. Ruiz-González, R., Cortajarena, A. L., Mejias, S. H., Agut, M., Nonell, S. & Flors, C. Singlet oxygen generation by the genetically encoded tag miniSOG. J. Am. Chem. Soc. 135, 9564–9567 (2013).

28. Riega, S. D. E., Rodríguez, H. B. & Román, E. S. Rose bengal in poly(2-hydroxyethyl methacrylate) thin films: self-quenching by photoactive energy traps. Methods Appl. Fluoresc. 5, 014010 (2017).

29. Gomes, A., Fernandes, E. & Lima, J. L. F. C. Fluorescence probes used for detection of reactive oxygen species. J. Biochem. Biophys. Methods 65, 45–80 (2005).

30. Bilski, P. J., Karriker, B. & Chignell, C. F. Quenching and generation of singlet oxygen by hydroethidine and related chromophores. Chem. Phys. Lett. 475, 116–119 (2009).

31. Wojtovich, A. P. & Foster, T. H. Optogenetic control of ROS production. Redox Biol 2, 368–376 (2014).

32. Liu, X. & Hajnóczky, G. Altered fusion dynamics underlie unique morphological changes in mitochondria during hypoxia–reoxygenation stress. Cell Death & Differentiation 18, 1561–1572 (2011).

33. Miyazono, Y., Hirashima, S., Ishihara, N., Kusukawa, J., Nakamura, K.-I. & Ohta, K. Uncoupled mitochondria quickly shorten along their long axis to form indented spheroids, instead of rings, in a fission-independent manner. Scientific Reports 8, 350 (2018).

34. Mitra, K., Wunder, C., Roysam, B., Lin, G. & Lippincott-Schwartz, J. A hyperfused mitochondrial state achieved at G1-S regulates cyclin E buildup and entry into S phase. Proc. Natl. Acad. Sci. U. S. A. 106, 11960–11965 (2009).

35. Tricot, S., Siberchicot, C., Bontemps, I., Desmaze, C., Kratassiouk, G., Vandamme, M., Pinna, G., Nemazanyy, I., Radicella, J. P., Lenaers, G., Renaudin, X., Lebeau, J. & Campalans, A. G-quadruplex ligand RHPS4 compromises cellular radioresistance by inhibiting the mitochondrial adaptive response induced by ionizing irradiation. NAR Cancer 7, zcaf033 (2025).

36. Hall, D. B., Holmlin, R. E. & Barton, J. K. Oxidative DNA damage through long-range electron transfer. Nature 382, 731–735 (1996).

37. Shibutani, S., Takeshita, M. & Grollman, A. P. Insertion of specific bases during DNA synthesis past the oxidation-damaged base 8-oxodG. Nature 349, 431–434 (1991).

38. Boiteux, S., Coste, F. & Castaing, B. Repair of 8-oxo-7,8-dihydroguanine in prokaryotic and eukaryotic cells: Properties and biological roles of the Fpg and OGG1 DNA N-glycosylases. Free Radic. Biol. Med. 107, 179–201 (2017).

39. Harada, K., Kubo, H., Abe, J., Haneta, M., Conception, A., Inoue, S., Okada, S. & Nishioka, K. Discovery of potent and orally bioavailable 17β-hydroxysteroid dehydrogenase type 3 inhibitors. Bioorg. Med. Chem. 20, 3242–3254 (2012).

40. Geldenhuys, W. J., Yonutas, H. M., Morris, D. L., Sullivan, P. G., Darvesh, A. S. & Leeper, T. C. Identification of small molecules that bind to the mitochondrial protein mitoNEET. Bioorg. Med. Chem. Lett. 26, 5350–5353 (2016).

41. Song, Y., Connor, D. T., Doubleday, R., Sorenson, R. J., Sercel, A. D., Unangst, P. C., Roth, B. D., Gilbertsen, R. B., Chan, K., Schrier, D. J., Guglietta, A., Bornemeier, D. A. & Dyer, R. D. Synthesis, structure-activity relationships, and in vivo evaluations of substituted di-tert-butylphenols as a novel class of potent, selective, and orally active cyclooxygenase-2 inhibitors. 1. Thiazolone and oxazolone series. J. Med. Chem. 42, 1151–1160 (1999).

42. Bottone, S., Broch, F., Gedeon, A., Brion, A., El Hajji, L., Benaissa, H. & Gautier, A. Design and evaluation of a tripartite chemogenetic fluorescent reporter for visualizing ternary protein complexes. Nat. Commun. 10.1038/s41467-025-62241-8 doi:10.1038/s41467-025-62241-8.

43. El Hajji, L., Benaissa, H. & Gautier, A. Isolating and engineering fluorescence-activating proteins using yeast surface display. Methods Mol. Biol. 2491, 593–626 (2022).

44. Adelizzi, B., Gielen, V., Le Saux, T., Dedecker, P. & Jullien, L. Quantitative model for reversibly photoswitchable sensors. ACS Sens. 6, 1157–1165 (2021).

45. Lahlou, A., Tehrani, H. S., Coghill, I., Shpinov, Y., Mandal, M., Plamont, M.-A., Aujard, I., Niu, Y., Nedbal, L., Lazár, D., Mahou, P., Supatto, W., Beaurepaire, E., Eisenmann, I., Desprat, N., Croquette, V., Jeanneret, R., Le Saux, T. & Jullien, L. Fluorescence to measure light intensity. Nat. Methods 20, 1930–1938 (2023).

