## Supplementary Information for "A chemogenetic light-activatable reactive oxygen species generator for precise cell manipulation"

[#] These authors contributed equally

This pdf file contains

Supplementary Figures 1 – 7

Supplementary Table 1

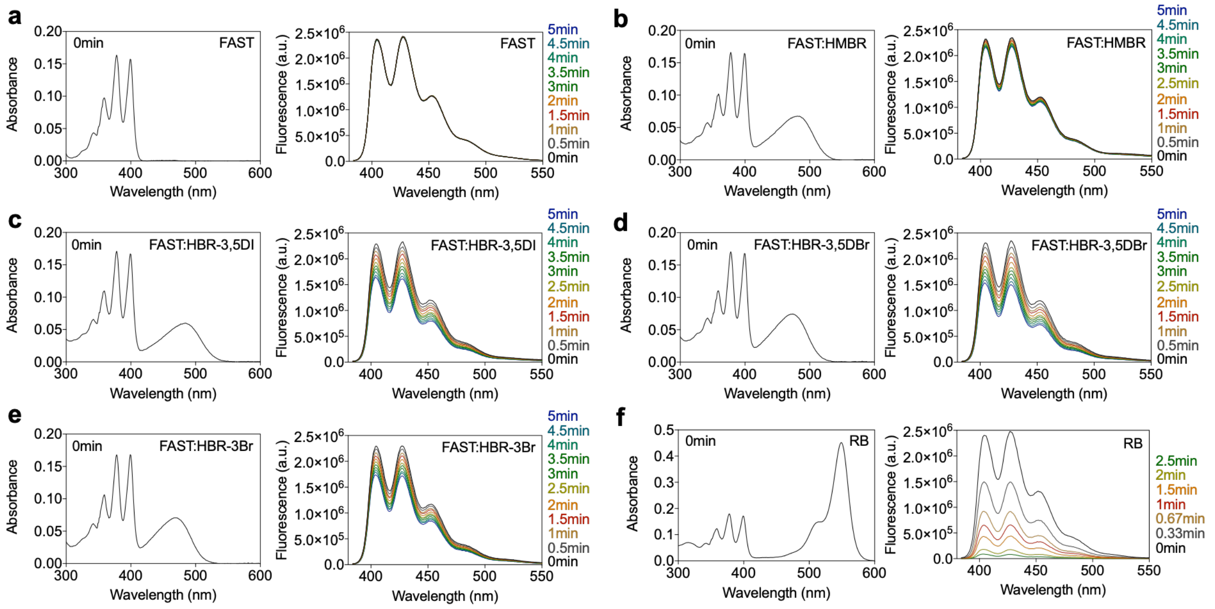


**Supplementary Figure 1. Anthracene-9-10-dipronoic acid (ADPA) photooxidation measurements in presence of a** FAST, **b** FAST:HMBR, **c** FAST:HBR-3,5DI, **d** FAST:HBR-3,5DBr, **e** FAST:HBR-3Br and **f** Rose Bengal (RB). Absorption (left) and fluorescence (right) spectra were recorded and ADPA photooxidation was monitored as a function of the duration of 488 nm light exposure (90 mW.cm$^{-2}$) (Ex 374 nm; Em 384 – 550 nm). Experiments were performed in presence of 50 μM ADPA and 15 μM of FAST; and the concentrations of HBR-3,5DI, HBR-3,5DBr, HBR-3Br (around 4 to 6 μM) and RB were adjusted to achieve optically matched solutions at 488 nm (abs$_0$ = 0.055 ± 10%).

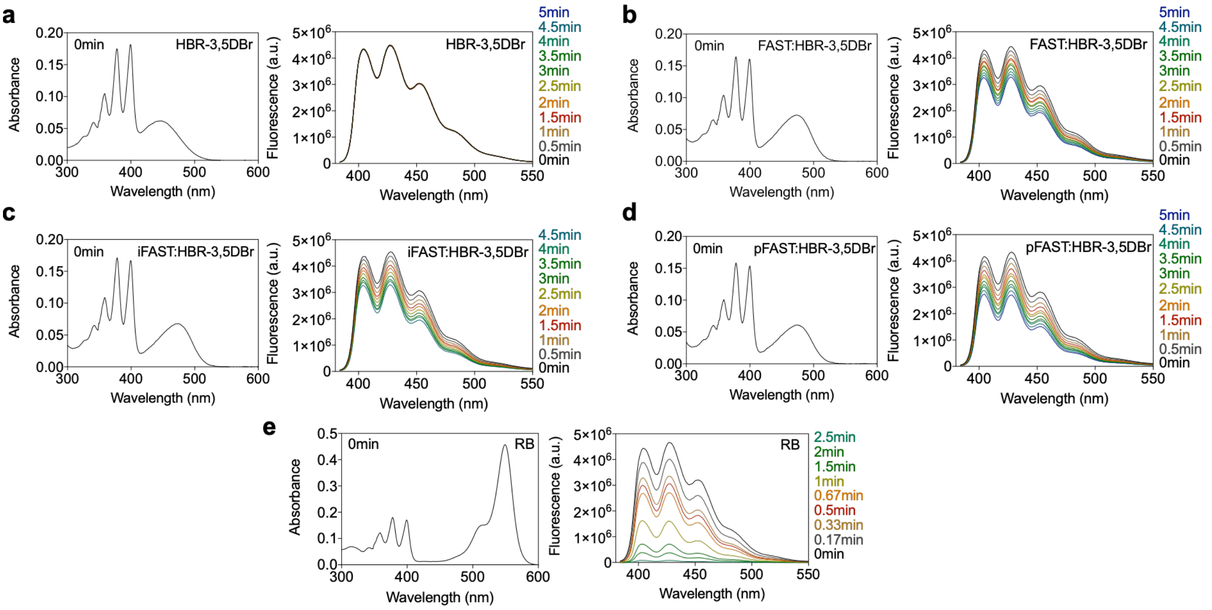


**Supplementary Figure 2. Anthracene-9-10-dipronoic acid (ADPA) photooxidation measurements in presence of a** HBR-3,5DBr, **b** FAST:HBR-3,5DBr, **c** iFAST:HBR-3,5DBr, **d** pFAST:HBR-3,5DBr, **e** Rose Bengal (RB). Absorption (left) and fluorescence (right) spectra were recorded and ADPA photooxidation was monitored as a function of the duration of 488 nm light exposure (90 mW.cm$^{-2}$) (Ex 374 nm; Em 384 – 550 nm emission). Experiments were performed in presence of 50 μM ADPA; FAST, iFAST and pFAST at 15 μM; and the concentrations of HBR-3,5DBr (4 – 5 μM), and RB were adjusted to achieve optically matched solutions at 488 nm ($abs_0$ = 0.055 ± 10%).

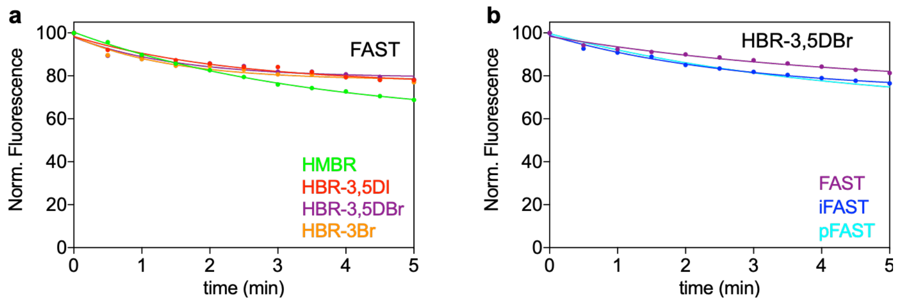


**Supplementary Figure 3. Photobleaching measurement of a** FAST (15 μM) in presence of HMBR, HBR-3,5DI, HBR-3,5DBr and HBR-3Br and **b** HBR-3,5DBr in presence of FAST, iFAST and pFAST at 15 μM. The concentration of chromophores (around 4 to 6 μM) was adjusted to achieve optically matched solutions at 488 nm ($abs_0$ = 0.055 ± 10%). Photobleaching was monitored under the same illumination conditions as for ADPA photobleaching and HE oxidation experiments (488 nm light exposure, 90 mW.cm$^{-2}$) (Ex 488 nm; Em 498 – 700 nm). Data were fitted using one phase exponential decay.

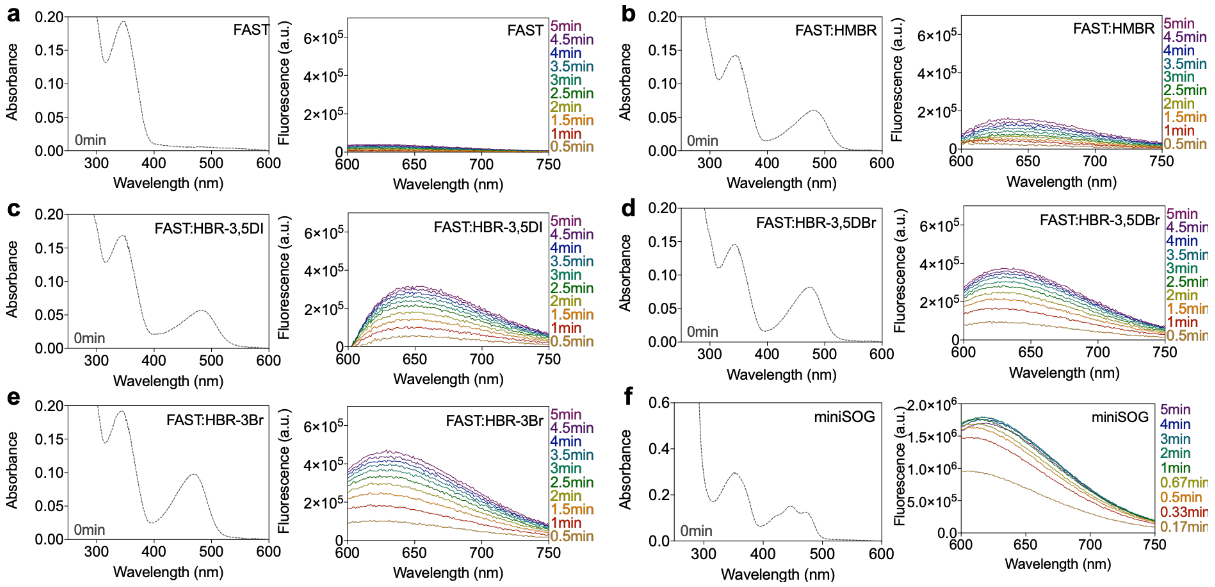


**Supplementary Figure 4. Dihydroethidium (HE) photooxidation measurements in presence of a** FAST, **b** FAST:HMBR, **c** FAST:HBR-3,5DI, **d** FAST:HBR-3,5DBr, **e** FAST:HBR-3Br and **f** miniSOG. Absorption (left) and fluorescence (right) spectra were recorded and HE oxidation was monitored as a function of the duration of 488 nm light exposure (90 mW.cm$^{-2}$) (Ex 525 nm; Em 600 – 750 nm). Experiments were performed in presence of 50 μM HE, 15 μM of FAST; and the concentrations of HBR-3,5DI, HBR-3,5DBr, HBR-3Br (around 4 to 6 μM) and miniSOG were adjusted to achieve optically matched solutions at 488 nm (abs$_0$ = 0.055 ± 10%).

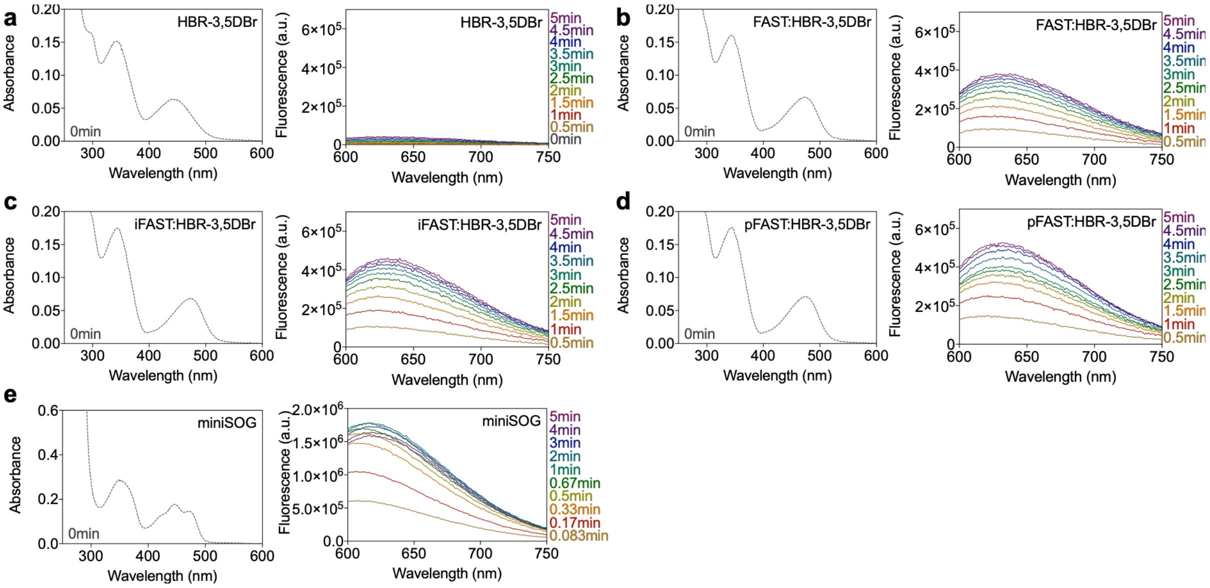


**Supplementary Figure 5. Dihydroethidium (HE) photooxidation measurements in presence of a** HBR-3,5DBr, **b** FAST:HBR-3,5DBr, **c** iFAST:HBR-3,5DBr, **d** pFAST:HBR-3,5DBr and **e** miniSOG. Absorption (left) and fluorescence (right) spectra were recorded and HE oxidation was monitored as a function of the duration of 488 nm light exposure (90 mW.cm$^{-2}$) (Ex 525 nm; Em 600 – 750 nm). Experiments were performed in presence of 50 μM HE; FAST, iFAST and pFAST at 15 μM and the concentrations of HBR-3,5DBr (4 – 5 μM) and miniSOG were adjusted to achieve optically matched solutions at 488 nm (abs$_0$ = 0.055 ± 10%).

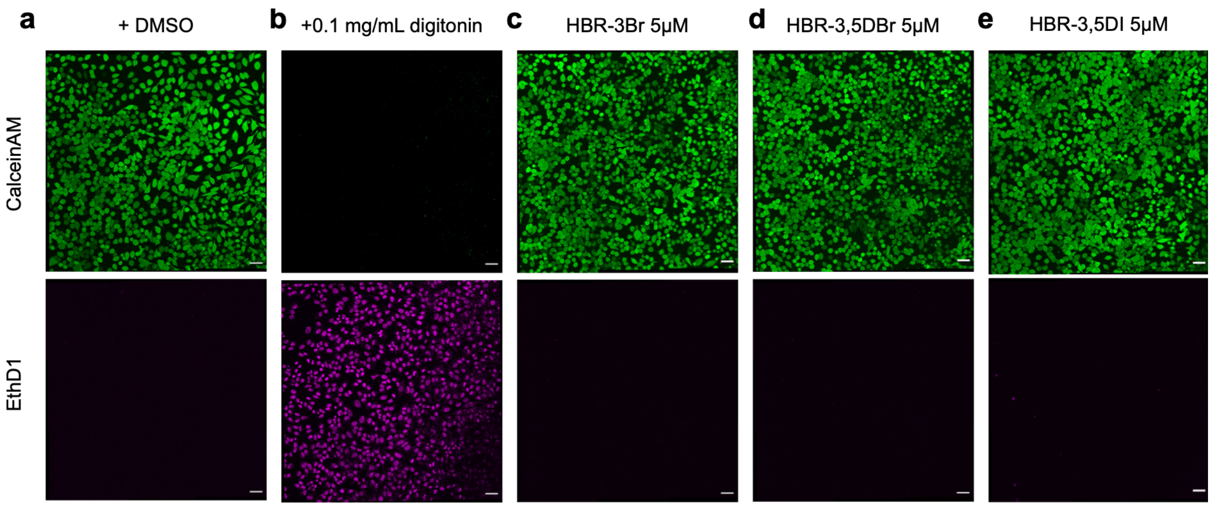


**Supplementary Figure 6. Two-color fluorescence viability assay.** HeLa cells were incubated for 24 h with solutions of **a,b** 0.1 % of DMSO, **c** HBR-3Br at 5 μM, **d** HBR-3,5DBr at 5 μM and **e** HBR-3,5DI at 5 μM. Control experiments of HeLa cells non-incubated with dye (**a**, live cells) or incubated for 30 min with 0.1 mg/ml digitonin (**b**, dead cells) are shown. Cell viability was tested with calceinAM and EthD1 probes (LIVE/DEAD® viability/cytotoxicity assay kit). CalceinAM is a cell-permeant profluorophore cleaved by intracellular esterases releasing a uniform green fluorescence polyanionic calcein in live cells (green channel). EthD1 (Ethidium homodimer-1) is a non-permeant nucleic acid fluorescent stain that enters only cells with damaged membranes and undergoes a fluorescence enhancement upon binding to nucleic acids, thereby producing a bright red fluorescence in dead cells (pink channel). Cell fluorescence was evaluated by confocal microscopy. Identical imaging settings were used for the different experiments. The experiment shows that none of the chromophores are toxic for HeLa cells at the concentrations used for labeling. Scale bars, 30 μm.

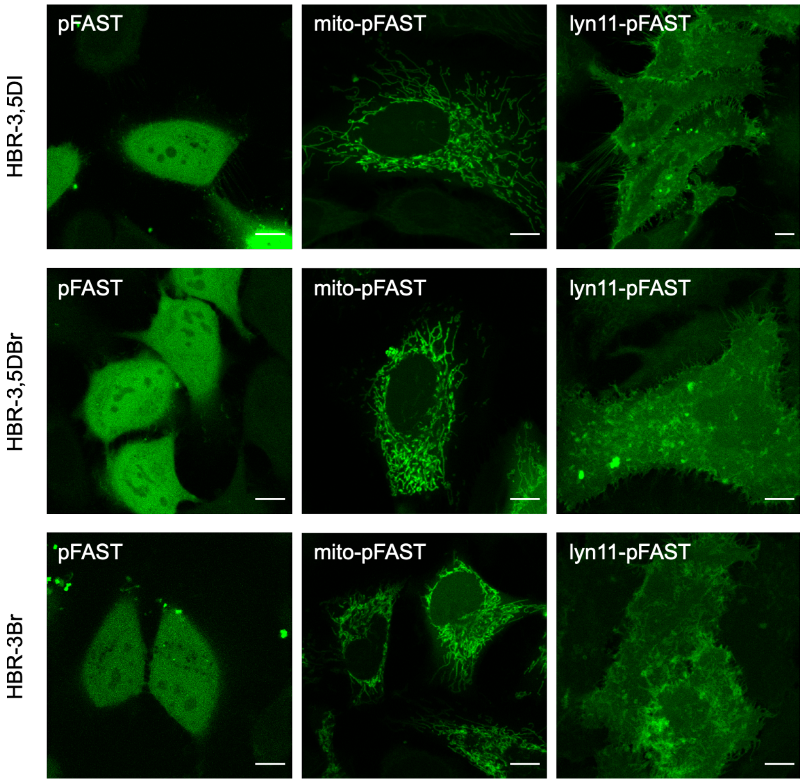


**Supplementary Figure 7. Selective imaging of pFAST in live mammalian cells**. Confocal micrographs of live HeLa cells expressing pFAST, or pFAST fused to the mitochondrial targeting sequence from the subunit VIII of human cytochrome C oxidase (Mito-pFAST) and to the inner plasma membrane targeting sequence lyn11 (Lyn11-pFAST) and labeled with 10 μM HBR-3,5DI, HBR-3,5DBr and HBR-3Br. Imaging conditions: Exc 488 nm, Em 500 – 590 nm. Scale bars, 10 μm.

**Supplementary Table 1. Plasmids used in this study.**

| Plasmid | Description | ORF sequence |
|---|---|---|
| pAG657 | H2B-pFAST-C-myc | **atg**cccgaacctgcgaagtcagcgcccgctcccaaaaaggctctaaaaagctgtcg<br>ccaagacccagaagaaggggataagaaaaggcgtaagaccaggaaagagagtta<br>cgccatttacgtgtacaaagtactaaaacaagtccacccggacactggcatctcctcaaa<br>ggcgatgggcattatgaactcatttgtaaacgacatcttcgagcgcatcgccggagaagc<br>gtcgcgcctggcgcattacaacaagcgctccactatcacatcccgggagatccagacgg<br>ccgtgcgcctgctcctgcccggagaactggccaaacacgctgtgtctgagggcacaaag<br>gccgtgaccaagtacaccagctccaagggcggaggctccggaggcggatctgccacc<br>atggagcatgttgcctttggcagtgaggacatcgagaacactctggccaatatggacgac<br>gaacaactggataggttggcctttggcgtaattcagctcgatggtgacgggaatatcctgct<br>gtacaatgctgctgaaggggacatcactggcagagatcccaaacaggtgattgggaag<br>aacttcttcaaggatgttgcacctggaacggatactcccgagttttacggcaaattcaagga<br>aggcgcagcgtcagggaatctgaacaccatgttcgaatggacgataccgacaagcagg<br>ggaccaaccaaggtcaaggtgcacttgaagaaagccctttccggtgacagatattgggtc<br>tttgtgaaacgggtgggatccgaacaaaagcttatttctgaagaggacttgtaa |
| pAG671 | Mito-pFAST-C-myc | **atg**tccgtcctgacgccgctgctgctgcggggcttgacaggctcggcccggcggctccca<br>gtgccgcgcgccaagatccattcgttgagatctgccaccatggagcatgttgcctttggcag<br>tgaggacatcgagaacactctggccaatatggacgacgaacaactggataggttggcctt<br>tggcgtaattcagctcgatggtgacgggaatatcctgctgtacaatgctgctgaaggggac<br>atcactggcagagatcccaaacaggtgattgggaagaacttcttcaaggatgttgcacctg<br>gaacggatactcccgagttttacggcaaattcaaggaaggcgcagcgtcagggaatctg<br>aacaccatgttcgaatggacgataccgacaagcaggggaccaaccaaggtcaaggtg<br>cacttgaagaaagccctttccggtgacagatattgggtctttgtgaaacgggtgggatccga<br>acaaaagcttatttctgaagaggacttgtaa |
| pAG1633 | mCherry-pFAST-(PH)2 | atggtgagcaagggcgaggaggataacatggccatcatcaaggagttcatgcgcttcaa<br>ggtgcacatggagggctccgtgaacggccacgagttcgagatcgagggcgagggcga<br>gggccgcccctacgagggcacccagaccgccaagctgaaggtgaccaagggtggcc<br>ccctgcccttcgcctgggacatcctgtcccctcagttcatgtacggctccaaggcctacgtg<br>aagcaccccgccgacatccccgactacttgaagctgtccttccccgagggcttcaagtgg<br>gagcgcgtgatgaacttcgaggacggcggcgtggtgaccgtgacccaggactcctccct<br>gcaggacggcgagttcatctacaaggtgaagctgcgcggcaccaacttcccctccgacg<br>gccccgtaatgcagaagaagaccatgggctgggaggcctcctccgagcggatgtaccc<br>cgaggacggcgccctgaagggcgagatcaagcagaggctgaagctgaaggacggcg<br>gccactacgacgctgaggtcaagaccacctacaaggccaagaagcccgtgcagctgc<br>ccggcgcctacaacgtcaacatcaagttggacatcacctcccacaacgaggactacacc<br>atcgtggaacagtacgaacgcgccgagggccgccactccaccggcggcatggacgag<br>ctgtacaagggcggcggctctggaggttccggaatggagcatgttgcctttggcagtgagg<br>acatcgagaacactctggccaatatggacgacgaacaactggataggttggcctttggcg<br>taattcagctcgatggtgacgggaatatcctgctgtacaatgctgctgaaggggacatcact<br>ggcagagatcccaaacaggtgattgggaagaacttcttcaaggatgttgcacctggaacg<br>gatactcccgagttttacggcaaattcaaggaaggcgcagcgtcagggaatctgaacac<br>catgttcgaatggacgataccgacaagcaggggaccaaccaaggtcaaggtgcacttg<br>aagaaagccctttccggtgacagatattgggtctttgtgaaacgggtgggatccggtggtg<br>gaagcggtggtggcatggactcgggccgggacttcctgaccctgcacggcctacaggat<br>gatgaggatctacaggcgctgctgaagggcagccagctcctgaaggtgaagtccagctc<br>atggaggagagagcgcttctacaagttgcaggaggactgcaagaccatctggcaggag<br>tcccgcaaggtcatgcggaccccggagtcccagctgttctccatcgaggacattcaggag<br>gtgcgaatggggcaccgcacggagggtctggagaagttcgcccgtgatgtgcccgagg<br>accgctgcttctccattgtcttcaaggaccagcgcaatacactagacctcatcgccccatcg<br>ccagctgatgcccagcactgggtgctggggctgcacaagatcatccaccactcaggctcc<br>atggaccagcgtcagaagctacagcactggattcactcctgcttgcgaaaagctgacaa<br>aaacaaggacaacaagatgagcttcaaggagctgcagaacttcctgaagggtggaggt<br>agtggcggtggaatggactcgggccgggacttcctgaccctgcacggcctacaggatga<br>tgaggatctacaggcgctgctgaagggcagccagctcctgaaggtgaagtccagctcat<br>ggaggagagagcgcttctacaagttgcaggaggactgcaagaccatctggcaggagtc<br>ccgcaaggtcatgcggaccccggagtcccagctgttctccatcgaggacattcaggaggt |

| | | |
|---|---|---|
| | | gcgaatggggcaccgcacggagggtctggagaagttcgcccgtgatgtgcccgaggac<br>cgctgcttctccattgtcttcaaggaccagcgcaatacactagacctcatcgccccatcgcc<br>agctgatgcccagcactgggtgctggggctgcacaagatcatccaccactcaggctccat<br>ggaccagcgtcagaagctacagcactggattcactcctgcttgcgaaaagctgacaaaa<br>acaaggacaacaagatgagcttcaaggagctgcagaacttcctgaagtag |
| pAG1634 | mCherry_(PH)$_2$ | atggtgagcaagggcgaggaggataacatggccatcatcaaggagttcatgcgcttcaa<br>ggtgcacatggagggctccgtgaacggccacgagttcgagatcgagggcgagggcga<br>gggccgcccctacgagggcacccagaccgccaagctgaaggtgaccaagggtggcc<br>ccctgcccttcgcctgggacatcctgtcccctcagttcatgtacggctccaaggcctacgtg<br>aagcaccccgccgacatccccgactacttgaagctgtccttccccgagggcttcaagtgg<br>gagcgcgtgatgaacttcgaggacggcggcgtggtgaccgtgacccaggactcctccct<br>gcaggacggcgagttcatctacaaggtgaagctgcgcggcaccaacttcccctccgacg<br>gccccgtaatgcagaagaagaccatgggctgggaggcctcctccgagcggatgtaccc<br>cgaggacggcgccctgaagggcgagatcaagcagaggctgaagctgaaggacggcg<br>gccactacgacgctgaggtcaagaccacctacaaggccaagaagcccgtgcagctgc<br>ccggcgcctacaacgtcaacatcaagttggacatcacctcccacaacgaggactacacc<br>atcgtggaacagtacgaacgcgccgagggccgccactccaccggcggcatggacgag<br>ctgtacaagggatccggtggtggaagcggtggtggcatggactcgggccgggacttcctg<br>accctgcacggcctacaggatgatgaggatctacaggcgctgctgaagggcagccagct<br>cctgaaggtgaagtccagctcatggaggagagagcgcttctacaagttgcaggaggact<br>gcaagaccatctggcaggagtcccgcaaggtcatgcggaccccggagtcccagctgttc<br>tccatcgaggacattcaggaggtgcgaatggggcaccgcacggagggtctggagaagtt<br>cgcccgtgatgtgcccgaggaccgctgcttctccattgtcttcaaggaccagcgcaataca<br>ctagacctcatcgccccatcgccagctgatgcccagcactgggtgctggggctgcacaag<br>atcatccaccactcaggctccatggaccagcgtcagaagctacagcactggattcactcct<br>gcttgcgaaaagctgacaaaaacaaggacaacaagatgagcttcaaggagctgcaga<br>acttcctgaagggcggcggctctggaggttccggaatggactcgggccgggacttcctga<br>ccctgcacggcctacaggatgatgaggatctacaggcgctgctgaagggcagccagctc<br>ctgaaggtgaagtccagctcatggaggagagagcgcttctacaagttgcaggaggactg<br>caagaccatctggcaggagtcccgcaaggtcatgcggaccccggagtcccagctgttct<br>ccatcgaggacattcaggaggtgcgaatggggcaccgcacggagggtctggagaagtt<br>cgcccgtgatgtgcccgaggaccgctgcttctccattgtcttcaaggaccagcgcaataca<br>ctagacctcatcgccccatcgccagctgatgcccagcactgggtgctggggctgcacaag<br>atcatccaccactcaggctccatggaccagcgtcagaagctacagcactggattcactcct<br>gcttgcgaaaagctgacaaaaacaaggacaacaagatgagcttcaaggagctgcaga<br>acttcctgaagtag |
